# Generation and characterization of a tamoxifen-inducible, Cre driver rat for transgene expression in microglia

**DOI:** 10.1101/2025.07.17.665447

**Authors:** Elliot J. Glotfelty, Lamarque M. Coke, Evan E. Hart, Cole Rivell, Amy Phan, Braxton D. Greer, Reinis Svarcbahs, Elizabeth Fielding, Julie Necarsulmer, Lowella V. Fortuno, Francois Vautier, Patricia J. Gearhart, Geoffrey Schoenbaum, Christopher T. Richie, Brandon K. Harvey

## Abstract

Microglia are the resident immune cells of the central nervous system (CNS) and display diverse functions under both physiological and pathological conditions. The past decade has seen burgeoning interest in microglia function, with a variety of transgenic tools developed for specific genetic manipulation of microglia in various injury, disease, and developmental models. Although the majority of models have been developed in mice, the ability to manipulate microglia in rats provides additional advantages to studying microglial function in the brain especially related to complex behavior. Using BAC transgenesis, our lab has created a transgenic rat (Cx3cr1-CreERT2) that expresses a tamoxifen inducible Cre recombinase (CreERT2) under control of the microglial/macrophage specific fractalkine C-X3-C Motif Chemokine Receptor 1 (*Cx3cr1*) promoter. In mice, CreERT2 and other transgenes have been expressed in microglia using the *Cx3cr1* promoter, however, this is the first demonstration in rats. Importantly, these rats exhibit similar cognitive behaviors compared to their wildtype (WT) controls. Microglial specificity of inducible Cre expression was confirmed by breeding the novel Cx3cr1-CreERT2^+/-^ rat with a previously reported double floxed inverse open reading frame (DIO)-mCherry^+/-^ reporter rat to show tamoxifen inducible mCherry expression that colocalizes with the microglial marker Iba1. In addition, we utilize flow cytometry to demonstrate time and Cre dependent differences in recombination of Cx3cr1^+^ cells in the spleen, peripheral blood, and brain at two- and eight-weeks post-tamoxifen treatment. Overall, we have created a novel transgenic rat model for researchers to employ in understanding microglial and peripheral immune cell function in rats.

## 1. Introduction

Microglia are the resident immune cells of the central nervous system (CNS). Originating from the fetal yolk-sac during embryonic development, microglia continue as a self-renewing population after they have infiltrated the brain (Chen et al., 2017; Ginhoux et al., 2010; Sheng et al., 2015). Although microglia maintain homeostasis within the healthy brain, they are a heterogenous population of cells that function in a context-dependent manner. Their functions are diverse and include induction of pro- and anti-inflammatory pathways in response to a CNS injury, the organization of scar tissue around spinal cord injury (Bellver-Landete et al., 2019), chronic inflammation after traumatic brain injury and disease (Glotfelty et al., 2019; Kopp et al., 2022; Younger et al., 2019), and release of cytokines during viral infection (Chen et al., 2019). Cross talk between microglia and neighboring neuronal cells leads to changes in both populations of cells (Marinelli et al., 2019). For example, neuronal factors can influence microglial phenotypes (Schafer et al., 2012; Stevens et al., 2007), and microglia are required for the pruning of synapses during normal development (Perez-Alcazar et al., 2014) and learning-related plasticity (Nie et al., 2018). Under certain conditions, microglia can alter animal behavior (Spangenberg et al., 2019) and are necessary for the initiation of plaque formation in the 5xFAD Alzheimer’s disease model (Veremeyko et al., 2019).

Over the past decade, several groups have manipulated microglia in rodents using viral vectors. These include the combination of novel “degradation-resistant” serotypes of Adeno-Associated Virus (AAV) with “microglia-specific” promoter elements (Rosario et al., 2016), VSV-G pseudo typed lentiviral vectors with ubiquitous promoters that were restricted to microglia expression using micro-RNA (miRNA) targeting elements (Åkerblom et al., 2013), and AAV vectors using portions of the microglia/macrophage-specific ionized calcium-binding adaptor molecule 1 (*Iba1*), along with miRNA target sites (Okada et al., 2022; Serrano et al., 2024). Although capable of expressing a transgene in microglia, these vectors also produce expression in non-microglial cells. Despite temporally and spatially confining transgene expression, these tools still lack cellular specificity for studies where transgene expression must be restricted to microglia.

Currently, mice are the predominant rodent model for genetically manipulating microglia. Through genomic manipulation of mouse embryos, transgenes such as fluorescent reporters or recombinases are expressed from “microglia-specific” promoters such as Cx3cr1 (Parkhurst et al., 2013; Yona et al., 2013), Tmem119 (Kaiser & Feng, 2019), P2ry12 (McKinsey et al., 2020), Sall1(Buttgereit et al., 2016), HexB (Masuda et al., 2020) and Csfr1 (Deng et al., 2010), with Cx3cr1 utilized most frequently. CX3CR1, also known as the fractalkine C-X3-C Motif Chemokine Receptor 1, is a seven transmembrane domain G-protein coupled receptor expressed in monocytes, natural killer (NK) and T cells, dendritic cells, and microglia (Jung et al., 2000), and is known to play a role in migration and adhesion of leukocytes (Imai et al., 1997), neuron-microglia interactions (Jung et al., 2000; Nishiyori et al., 1998), maintenance of synapses (Paolicelli et al., 2011), and in Human Immunodeficiency Virus (HIV) associated pathologies (Garin et al., 2003). Mouse models utilizing the Cx3cr1 promoter, among others, to drive microglial- specific transgenes, including Cre-recombinase and fluorescent reporters, have been used in neuroscience research to study microglial biology with varying degrees of specificity (Bedolla et al., 2024; Jung et al., 2000).

Mice are the primary mammalian animal model for human disease, due in part to the relatively low cost to maintain large numbers compared to other mammals, an extensive list of available transgenic lines, a fully sequenced and annotated genome, and tools to manipulate that genome. Rats, however, offer advantages over mice, including larger size for ease of surgical manipulation, more biomass to analyze, and behavioral complexity for physiology, toxicology, and neuroscience studies (Aitman et al., 2016). Given that microglia are essential for maintaining and restoring brain homeostasis under normal and pathological conditions (Paolicelli et al., 2022) and affecting animal behavior (Chen et al., 2024), we sought to develop a microglia-specific genetic model in rats to expand our capabilities to study the role of microglia in complex behavioral paradigms such as addiction or neurodegeneration. Here, we describe a novel rat model for studying microglia based on a transgenic line expressing the tamoxifen-inducible CreERT2 recombinase under the control of the rat Cx3cr1 promoter. We characterize this rat on both Long Evans (LE) and Sprague-Dawley (SD) backgrounds and demonstrate inducibility and specificity of transgene expression in peripheral immune cells and microglia in the rat brain.

## Results

### 2.1 Creation of Cx3cr1-CreERT2 and microglial specific tamoxifen inducible DIO-mCherry rats

We first created an AAV vector with a transgene cassette containing the CreERT2 coding region (Metzger & Chambon, 2001) and the polyadenylation signal from bovine growth hormone (pAAV EF1α CreERT2; **Fig. S1A**). This construct was functionally validated for tamoxifen-dependent recombinase activity by packaging into AAV1 particles and transducing rat primary cortical neurons (PCNs) along with a Cre- dependent mCherry reporter virus AAV EF1α DIO mCherry, (Richie et al., 2017). Minimal mCherry expression was observed in vehicle treated cells, and the addition of 1µM 4-hydroxytamoxifen (4-OHT) with AAV EF1α CreERT2 or AAV EF1α iCre increased mCherry expression in primary cultures (**Fig. S1B and C**). After confirming the CreERT2 cassette was capable of promoting recombination, it was used to replace the start codon of the Cx3cr1 coding region in a bacterial artificial chromosome (BAC), carrying a 137 kilobase segment of the rat genome containing the *Cx3cr1* gene [**Fig. 1A**, (Tischer et al., 2010)]. The resulting Cx3cr1-CreERT2 BAC was injected into embryos isolated from (LE) rats to create founder rats “LE-Tg(Cx3cr1-CreERT2)3Ottc”. The LE-Tg(DIO-mCherry)2Ottc rat (RRRC ID # 00687) (Bäck, Necarsulmer, et al., 2019)was created by embryonic injection of plasmid with an EF1α promoter driving expression of double-floxed mCherry cassette (**Fig. 1B**). The DIO-mCherry^+/-^ rat was crossed with the Cx3cr1-CreERT2^+/-^ rat to examine the specificity of Cre expression control in microglia (**Fig. 1C**).

**Figure 1:**
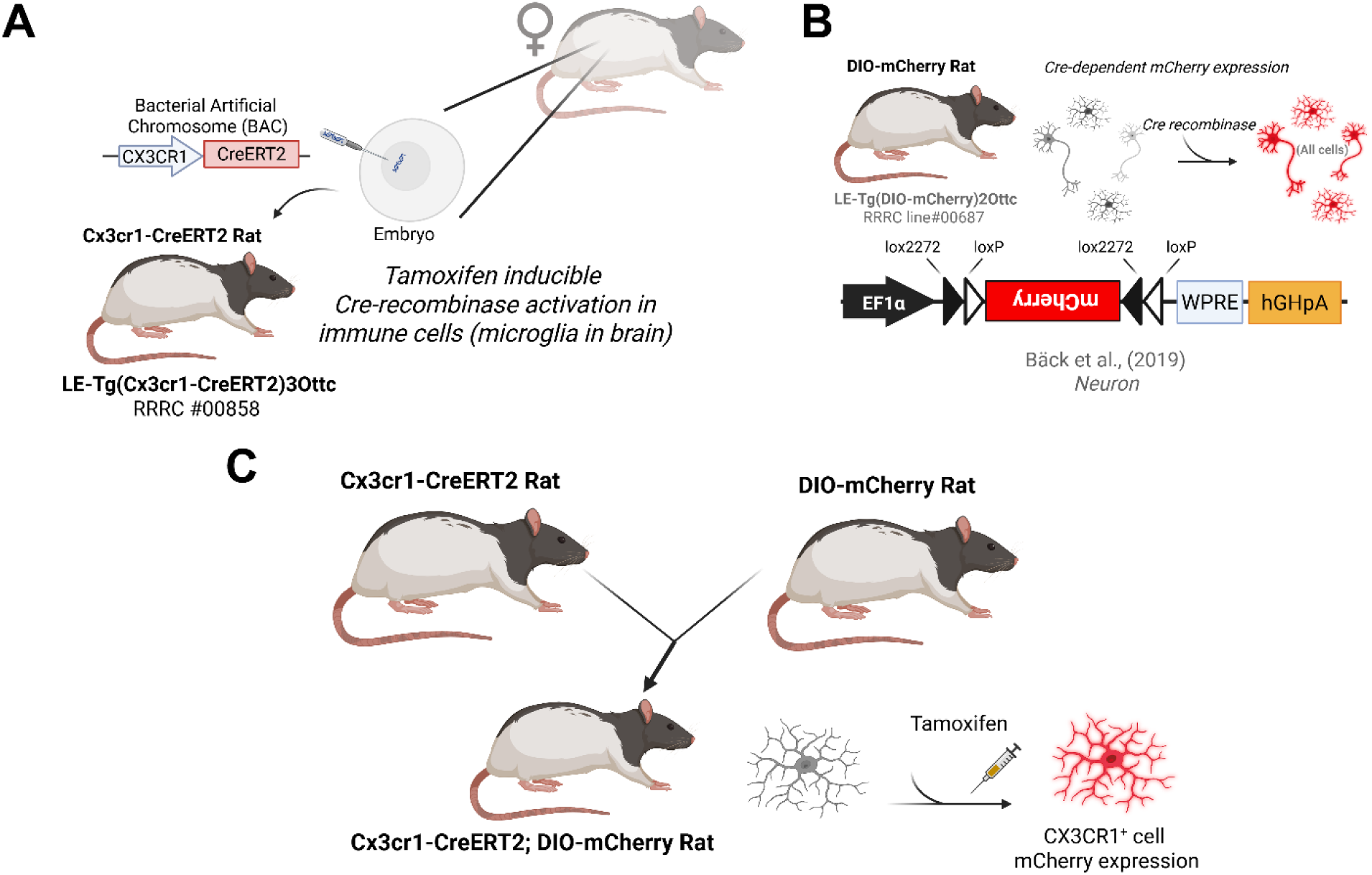
Cx3cr1-CreERT2 rat creation and characterization overview. **(A)** The Cx3cr1-CreERT2 DNA was injected into Long Evans (LE) rat embryos. **(B)** The double-floxed inverted open reading frame (DIO) was used to produce the DIO-mCherry rat for Cre-dependent expression of the mCherry fluorescent protein in any cells that express Cre. **(C)** The DIO-mCherry rat was crossed with the Cx3cr1-CreERT2 and treated with tamoxifen (TAM) to fluorescently label Cx3cr1 expressing cells such as microglia in the brain.

### 2.2 Genetic and behavioral characterization of the Cx3cr1-CreERT2 rat

To confirm effectiveness of BAC transgenesis and integration of the Cx3cr1-CreERT2 transgene, we isolated genomic DNA from tissue samples from pups of Cx3cr1-CreERT2^+/-^ × Cx3cr1-CreERT2^+/-^ crosses (**Fig. 2A**). We used 5’ and 3’ junction PCR and observed that (Hem) and homozygous (Hom) animals show identical 5’ and 3’ junction lengths, while wildtype (WT) animals exhibit no amplification of either transgene junction (**Fig. 2B**). We used ddPCR to quantify copies of integrated Cx3cr1-CreERT2 transgenes compared to the single copy gene Ggt1 (Kurauchi et al., 1991). We determined that Hem and Hom animals have 10 and 20 detectable copies of the transgene per diploid genome, respectively. No copies are expressed in the WT animals (**Fig. 2C**). Importantly, we confirm with digital PCR (dPCR) that mRNA expression of Cx3cr1 native locus is comparable in the WT and transgenic animals (**Fig. 2D**), while CreERT2 transgene is expressed in Hem animals and absent in the WT group (**Fig. 2E**).

**Figure 2:**
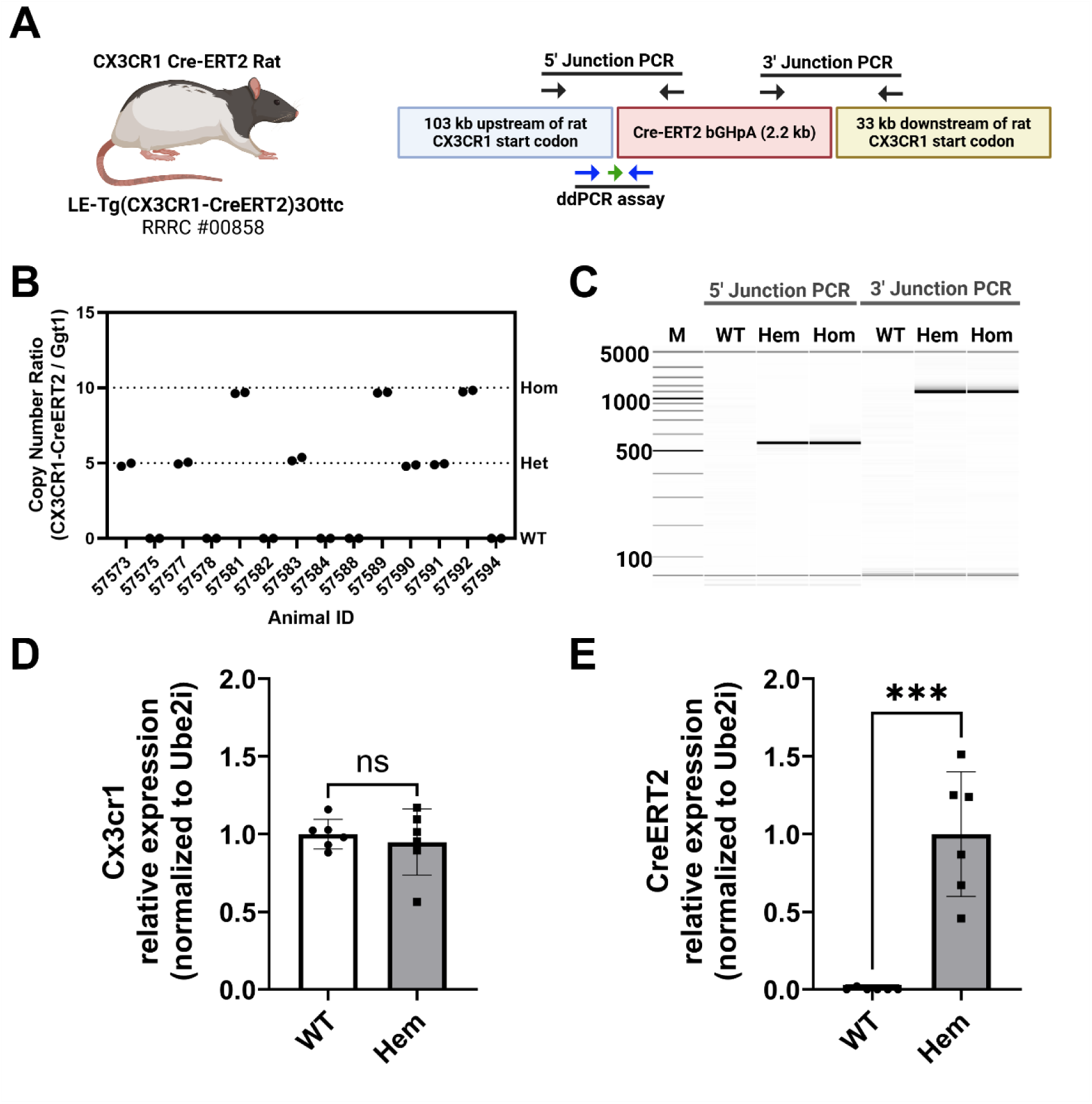
Cx3cr1-CreERT2 rat genetic characterization. **(A)** The Cx3cr1-CreERT2 rat was created from a bacterial artificial chromosome (BAC) containing the rat Cx3cr1 gene (CHORI) and recombineered to with the coding region of CreERT2 and the poly-adenylation signal of bovine growth hormone (bGHpA) immediately after the start codon of the rat Cx3cr1 gene. The primer positions (black arrows) for PCR genotyping of the 5’ and 3’ junctions of the CreERT2 DNA cassette. The positions of the primers (blue arrows) and probe (green arrow) for a droplet digital PCR (ddPCR)- and digital PCR (dPCR)- based genotyping assay/gene expression assays are shown along the bottom. **(B)** Digital gel image showing PCR results from the 5’ and 3’ junction assays in non-carrier animals (WT), hemizygous (Hem), and homozygous (Hom) carriers of the Cx3cr1-CreERT2 transgene. The 5’ and 3’ junction amplicons are 550 bp and 1553 bp, respectively. **(C)** ddPCR assay results from genomic DNA isolated from hemizygous and homozygous animals. Relative RNA expression of ectopic Cx3cr1 locus **(D)** and Cx3cr1-CreERT2 locus in WT and transgenic animals using dPCR. As expected, only transgenic animals express the CreERT2 transgene (t-test p<.001).

To ensure the presence of copies of the Cx3cr1 transgene does not disrupt behavioral phenotypes, we measured rats’ ability to perform a standard sensory preconditioning task (Brogden, 1939; Hart et al., 2020) (**Fig. 3A**). This occurred in three stages: in the first stage (preconditioning) rats were presented with two serial pairs of sensory cues, and in the second (reward conditioning) one cue was paired with rewards while the second served as the control cue. During the final stage (probe test), rats were presented with each of the four sensory cues, without rewards. We measured the percent time spent in the food port minus baseline responding (CS-baseline) during each stage. Higher levels of food port responding indicate reward-expectation and, therefore, learning about the cues (Delamater, 1995). If rats learned the initial cue pairings during preconditioning, they would respond more to the cue that predicted the reward-paired cue than the control cue during the probe test, indicating their ability to make inferences is intact (Sadacca et al., 2018; Wang, Howard, et al., 2020; Wang, Schoenbaum, et al., 2020) .

**Figure 3:**
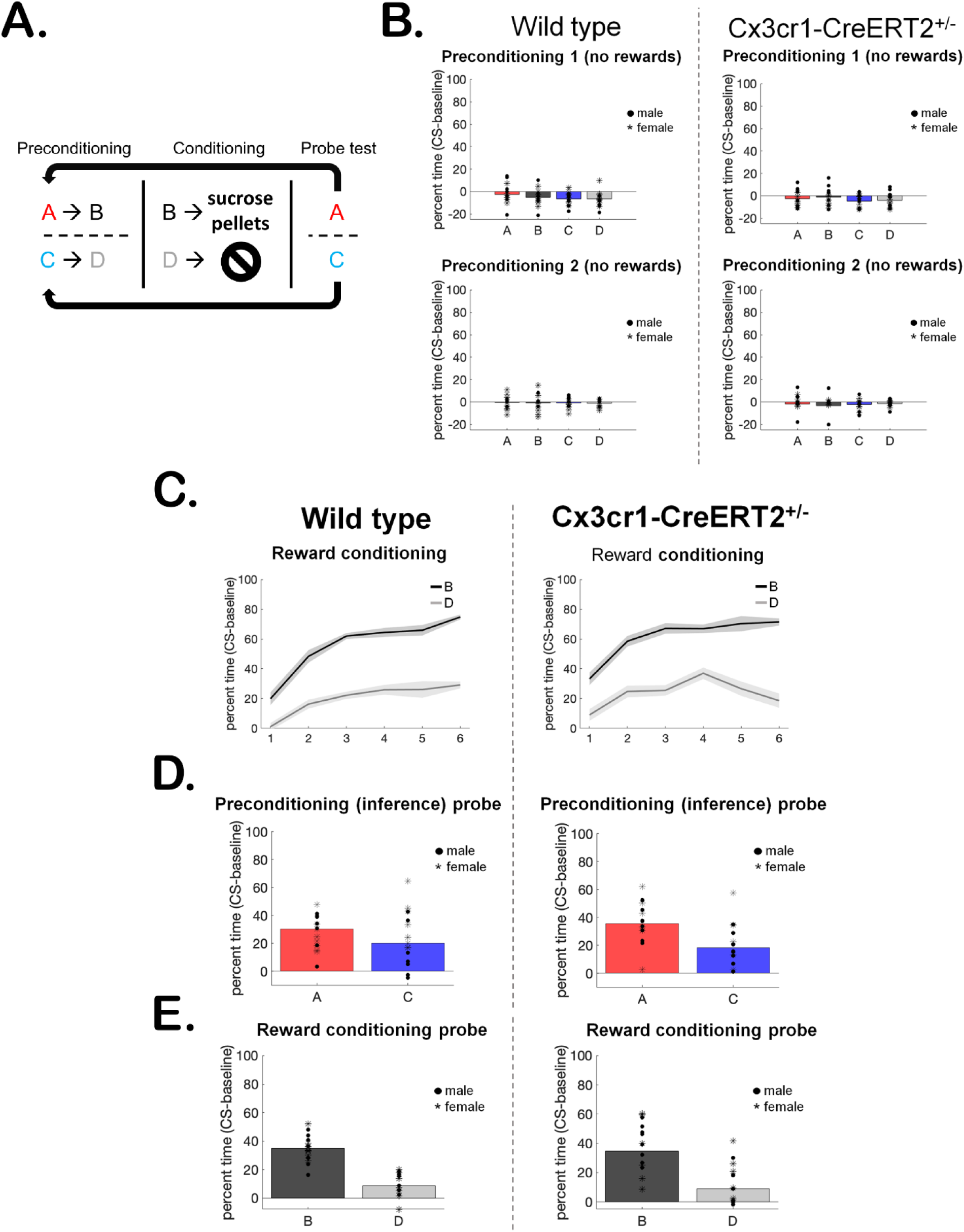
Behavior of LE Cx3cr1-CreERT2^+/-^ rats. **(A)** Sensory preconditioning behavioral task schematic. **(B)** WT and Hem rats responded similarly and at low levels to the neutral sensory cues during both days of preconditioning. **(C)** WT and Hem rats exhibited similar reward-learning during the six conditioning sessions, indicated by higher responding to the reward-paired cue B than to control cue D. **(D)** WT and Hem rats exhibited similar inference behavior during the probe test, indicated by higher responding to the cue that predicted the reward-paired cue during preconditioning (A>C responding). **(E)** WT and Hem rats exhibited similar reward learning and memory during the probe test, indicated by higher responding to the reward-paired cue B than to control cue D.

As expected, rats did not respond to the neutral cue pairs during the first or second day of preconditioning. WT and Hem rats responded similarly (**Fig. 3B**, all *p*-values >0.2, full table displayed in **Table 1**), since no rewards were involved. We next assessed whether rats could perform standard reward learning. Again, as expected, both groups learned to respond more to the reward-paired cue B than to the control cue D (**Fig. 3C**, main effect of session on B versus D responding, all other *p*-values >0.2). Finally, we tested whether WT and Hem rats exhibited similar inference behavior, which would be indicated by higher responding to the cue that predicted the reward-paired cue (cue A) than to the cue that predicted the control cue (cue C). During this final probe test, when no rewards were presented, rats in both groups exhibited typical inference behavior (**Fig. 3D**), responding more to cue A than to cue C (main effect of cue, all other *p*-values >0.07). Reward learning and memory were similarly intact, indicated by higher responding to cue B than cue D (**Fig. 3E**, main effect of cue, all other *p*-values >0.08). In sum, our behavioral results indicate that the Cx3cr1-CreERT2 transgene did not interfere with rats’ ability to form associative memories and make inferences based on them. A full report on behavioral statistics is displayed in **Table 1**.

**Table 1.** Behavioral statistics. Significant *p*-values bolded.

|  | Main effect<br>- cue | Main effect<br>- group<br>(WT vs<br>Hem) | Main effect<br>- sex | Main effect<br>- session | 2-way<br>interaction<br>(group x cue) | 3-way<br>interaction<br>(group x sex x<br>cue) |
| --- | --- | --- | --- | --- | --- | --- |
| Preconditioning 1 | F(3,75)=2.98,<br>p=0.10 | F(1,25)=0.77,<br>p=0.39 | F(1,25)=0.38,<br>p=0.54 | - | F(3,75)=0.89, p=0.36 | F(3,75)=1.02,<br>p=0.32 |
| Preconditioning 2 | F(3,75)=0.22,<br>p=0.65 | F(1,25)=0.67,<br>p=0.42 | F(1,25)=1.52,<br>p=0.23 | - | F(3,75)=0.40, p=0.53 | F(3,75)=1.06,<br>p=0.31 |
| Reward<br>conditioning (B<br>versus<br>D<br>responding) | - | F(1,25)=0.15,<br>p=0.71 | F(1,25)=0.41,<br>p=0.53 | F(5,125)=9.<br>95, **p=0.004** | F(5,125)=0.83,<br>p=0.37 | F(5,125)=1.27,<br>p=0.27 |
| Probe test – A versus<br>C, inference | F(1,25)=4.72,<br>**p=0.04** | F(1,25)=0.87,<br>p=0.36 | F(1,25)=3.04,<br>p=0.09 | - | F(1,25)=1.37, p=0.25 | F(1,25)=0.74,<br>p=0.40 |
| Probe test – B versus<br>D, reward learning | F(1,25)=94.37<br>, **p<0.0001** | F(1,25)=0.72,<br>p=0.40 | F(1,25)=0.01,<br>p=0.92 | - | F(1,25)=0.01, p=0.91 | F(1,25)=3.27,<br>p=0.08 |

### 2.3 Characterization of the DIO-mCherry rats using AAV-Cre

We previously described the creation of the DIO-mCherry rat [(Bäck, Necarsulmer, et al., 2019); **Fig.4A**]. Here, we further characterized the expression of mCherry rats from both hemizygous (Hem) and Hom animals. The transgene construct and approximate PCR assay locations are shown in **Fig.4A**. We first confirmed zygosity by ddPCR normalizing to a genomic target (Ggt1). The individual ratios of templates (DIO-mCherry/Ggt1) fall into three quanta (0,0.5, and 1.0), indicating that the allele consists of a single copy of the transgene integrated into the genome (**Fig 4B**). Using standard ddPCR assay across the 5’ DIO region we detect the presence of transgene, but a qPCR assay is needed for determining zygosity depending on the breeding strategy used in additional animals (**Fig 4C**). Hem and Hom animals were identified and injected intrastriatally with an AAV-iCre expressing virus and were evaluated for ectopic mCherry and Cre expression after two weeks of recovery (**Fig. 4D**). mCherry expression is limited to areas proximal to AAV-iCre injections, shown in the upper panel of **Fig. 4D**. We did not evaluate which cell types were transduced, but independent of cell type, Cre expression from the AAV delivery correlates with mCherry expression [**Fig. 4D**, bottom panel, Overlap Coefficient (OC)=0.90]. In WT littermates, no mCherry expression was observed in AAV-iCre transduced cells (**Fig. S2A**). Hem and Hom DIO-mCherry rats exhibit similar AAV-iCre transduction (**. S2 B and C**), showing no difference in iCre immunostaining (**Fig. S2D**). Densitometry analysis of mCherry expression revealed a significant elevation in the integrated density of mCherry epifluorescence of Hom v. Hem animals (p<0.01) (**Fig. S2 E**).

**Figure 4:**
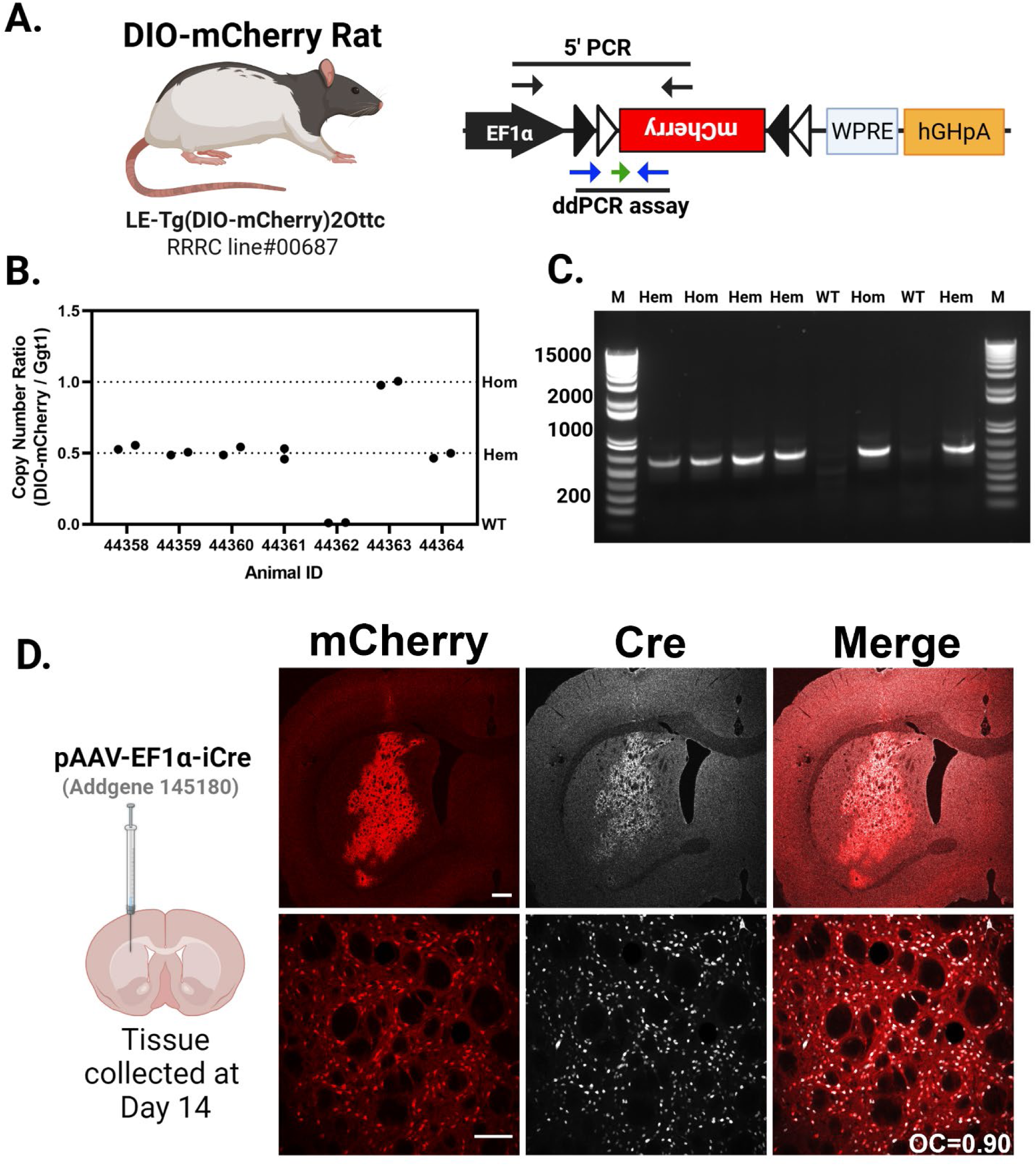
Cre-dependent mCherry rat. **(A)** The DIO-mCherry rat uses human EF1α promoter to drive expression of mCherry fluorescent protein along with the woodchuck post transcriptional regulatory element (WPRE) and the human growth hormone polyadenylation signal (hGHpA). **(A)** The primer positions for a PCR genotyping assay spanning the 5’ junction of the mCherry cassette is noted by the black arrows and the positions of the primers (blue) and probe (green arrow) for a qPCR-based genotyping assay are also noted. (**B**) Copy number for WT, hemizygous (Hem), and homozygous (Hom) animals. The ratio is transgene copy number to the genomic gene (Ggt1) in genomic DNA samples isolated from the pups of two separate Hem × Hem crosses. The individual ratios of templates (DIO-mCherry/Ggt1) fall into three quanta (0, 0.5, and 1.0). **(C)** Amplicons (640 bp) produced by the 5’ junction PCR identify wild type (no bands) or carriers hemizygous and homozygous. **(D)** Images from the striatum of hemizygous DIO-mCherry rats injected with an AAV virus expressing Cre recombinase after two weeks. Low magnification (top row; scale bar: 500 μm) and higher magnification (bottom row; scale bar: 50 μm). Cre immunoreactivity (white) and mCherry epifluorescence (red) colocalize in both sets of images. Overlap Coefficient (OC)=0.90.

### 2.4 Tamoxifen-induced Cre expression in Cx3cr1-CreERT2 rat brain colocalizes with Iba1^+^ cells

To further validate the Cx3cr1-CreERT2 construct, Cx3cr1-CreERT2^+/-^ rats were crossed with DIO-mCherry^+/-^ rats. Animals were intraperitoneally injected with tamoxifen (TAM) or vehicle (VEH) to activate the recombinase from postnatal day 10-12, with brains harvested two to eight weeks later (**Fig. 5A**). Immunofluorescence was performed with anti-mCherry antibody to provide more sensitive confirmation of reporter expression, along with the anti-Iba1 antibody to identify microglia. mCherry^+^Iba1^+^ cells were manually counted in four different brain regions [Pre-frontal cortex (PFC), Striatum (STR), Hippocampus (HC), and Midbrain (MB)] to evaluate colocalization. (**Fig. S3**.) In tandem, cells positive for anti-mCherry (antibody) immunofluorescence were counted in the same images to correlate with ectopic mCherry signal. Data appearing in **Figure 5** is associated with the animals 7-8 weeks post TAM (60mg/kg) or vehicle injections. In vehicle injected animals, minimal tamoxifen independent recombination was observed in each brain region (**Fig. 5B**), while TAM-mediated Cre activation increased mCherry expression in Iba1^+^ cells (**Fig. 5C**). In addition to Iba1^+^mCherry^+^ cells, we also observed small amounts of Iba1^+^mCherry^-^ and Iba1^-^mCherry^+^ cells (**Fig. 5D**). In the PFC, STR, HC, and MB of vehicle and TAM injected animals, we observed regional averages of 10-15% and 40-50% of mCherry^+^ cells colocalizing with Iba1^+^ cells and saw similar expression patterns when using the anti-mCherry antibody colocalization in vehicle (**Fig. S4 A**) and TAM treated animals (**Fig. S4 B-C**). Notably, two animals did not robustly respond to TAM treatment, bringing all averages down and producing wider deviations within groups. TAM dependent expression of mCherry in the brain (microglia) appears stable from 2-8 weeks with average percentage of Iba1^+^ cells expressing the transgene ranging from 54-67% two weeks post-TAM (**Fig. S5 A**) and 57-60.5% four weeks post-TAM (**Fig. S5 B**). In vehicle treated animals, mCherry expression in Iba^+^ cells range from 18-26% at two weeks post-TAM (**Fig. S5 A**) and 9-19.5% four weeks post-TAM (**Fig. S5 B**). At all timepoints (2, 4, and 7-8 weeks), TAM treatment significantly elevated mCherry expression in Iba1^+^ cells.

**Figure 5:**
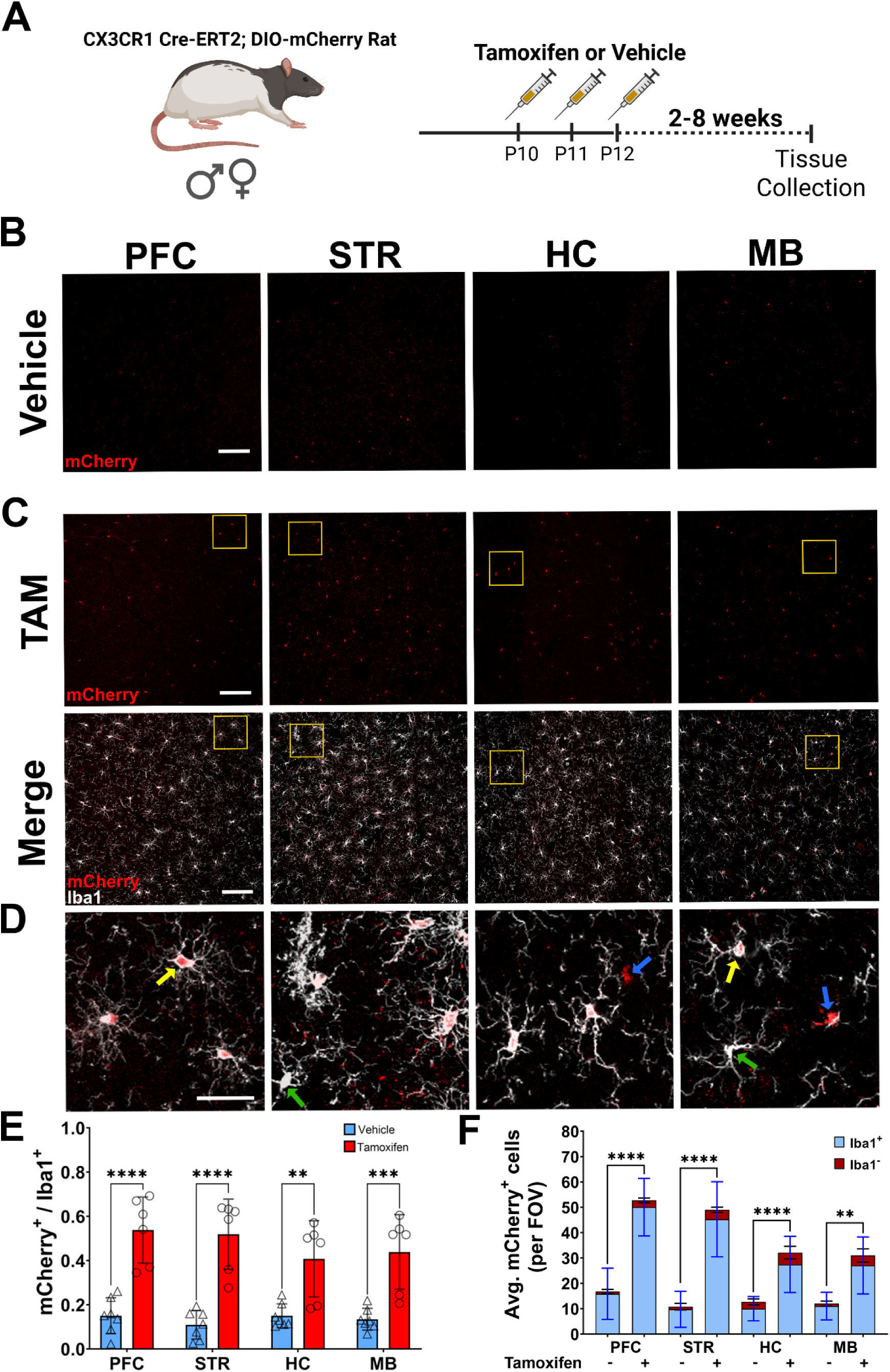
Specificity of Cre activity in microglia. **(A)** Male and female double transgenic animals (Cx3cr1-CreERT2^+/-^; DIO-mCherry^+/-^) received three i.p. injections of either tamoxifen (TAM) (male, n=3; female, n=4) or vehicle (male, n=4; female, n= 3) at P10. Brain tissue was collected 8 weeks after injections. Brains were sectioned and images taken proximal to the prefrontal cortex (PFC), striatum (STR), hippocampus (HC), and midbrain (MB) sections. Representative regional images show ectopic mCherry (red) expression in vehicle **(B)** and TAM **(C)** injected rats (scale bar:100 µm). Colocalization of ectopic mCherry (red) and Iba1 (white) in TAM injected rats is shown in the bottom panel of **(C)**. Regions shown at higher magnification in (**D)** (scale bar= 30 μm) are highlighted in yellow boxes from **(C)**. Yellow arrows indicate representative Iba1^+^ mCherry^+^ cells; green arrows highlight Iba1^+^mCherry^-^ cells; blue arrows show Iba1^-^ mCherry^+^ cells. **(E)** Ratio of mCherry^+^ /Iba1^+^ cells in vehicle (n=7) and TAM (n=6) treated rats (male and female animals combined). **(F)** Raw counts of mCherry^+^ cells in vehicle and TAM treated animals. Each bar indicates average quantity of mCherry^+^ cells that are Iba1^+^ (blue) and Iba1^-^ (red) per field of view. Two-way ANOVA with Tukey’s multiple comparison with S.D. error bars (**p<0.01; ***p<0.001; ****p<.0001).

In addition to the Long Evans background, the Cx3cr1-CreERT2 was backcrossed onto a SD background for at least 10 generations. We bred the SD animals with the DIO-mCherry rats (also on SD background), treated offspring with TAM (60mg/kg), and assessed, via immunohistochemistry, brain patterns of ectopic mCherry (red) and anti-mCherry (green) expression in Iba1^+^ (white) cells six weeks after treatment (**Fig. S6 A**). In the PFC, STR, HC, and MB, ectopic mCherry expression is restricted to Iba1^+^ cells, with similar anti-mCherry antibody colocalization (**Fig. S6 B**). Primary microglia were plated for *in vitro* evaluation of individual SD pups. Cells plated from Cx3cr1-CreERT2 negative pups showed almost no mCherry transgene expression when treated with vehicle (**Fig. S7 A**) or 4-hydroxytamoxifen (**Fig. S7 B**). Primary microglia from Cx3cr1-CreERT2-positive pups exhibit some mCherry expression in Iba1^+^ cells treated with vehicle (**Fig. S7 A**), as in our *ex vivo* studies. The level of mCherry expression in Iba1^+^ cells treated with 4-hydroxytamoxifen (4-OHT) increased compared to the vehicle treatment (**Fig. S7 B**).

### 2.5 Time and tissue dependent mCherry expression in LE Cx3cr1- CreERT2^+/-^; DIO-mCherry^+/-^ rats

Although the primary focus of the current studies evaluates brain specific expression of the Cx3cr1- CreERT2 transgene and mCherry reporter, Cx3cr1 is also expressed in a variety of peripheral cells including monocytes, dendritic cells, differentiating T-cells, and NK cells (Imai et al., 1997). Turnover rate of Cx3cr1^+^ peripheral circulating and immune cells is much faster (∼30 days) (Goldmann et al., 2013; Parkhurst et al., 2013; van Furth & Cohn, 1968; Yona et al., 2013) than the self-renewing population of microglia (Ajami et al., 2007) in the brain (∼95 days) (Askew et al., 2017). Peripheral cell recombination is a vital consideration when using the Cx3cr1-promoter driven transgene translation to assess microglia. In our rat model, we sought to explore expression of mCherry in the brain and periphery via flow cytometry (FC). In addition to assessing mCherry expression in Cx3cr1^+^ cells in the brain, we targeted the spleen and blood, as these tissues are reservoirs for Cx3cr1 expressing cells (Morgan et al., 2024; Yamauchi et al., 2021). We evaluated brain, blood, and spleen samples at 2- and 7-8-weeks post TAM treatment to understand recombination efficiency and stability of transgene expression over time.

We utilized a Cx3cr1 antibody to label cells of interest in the brain, blood, and spleen and then assessed for ectopic mCherry expression with and without TAM treatment (7-8 week animals). Experimental overview is shown in **Fig. 6A**. Representative gating charts of brain tissue 7-8 weeks after vehicle (top panel) and TAM (bottom panel) treatment demonstrates mCherry expression differences in Cx3cr1^+^ cells (**Fig 6B**). Representative gating of 7–8-week spleen and peripheral blood for vehicle and TAM treatments available in **Fig. S8 A** and **B**, respectively. Significant elevation (p=0.0061) in double positive (Q2) cells in TAM treated rats is observed at the 7–8-week time point (**Fig. 6C**). We observe no differences in mCherry expression in Cx3cr1^+^ cells in the peripheral blood (**Fig. 6D**) or spleens (**Fig. 6E**), which was expected given the turnover rates of recombined peripheral cells occurs much earlier than 7-8 weeks after TAM treatment. Expression of mCherry in Cx3cr1^+^ cells of the brain is stable between two and eight weeks as no significant differences were observed (**Fig. 6F**), Post TAM treatment, significant elevation of Cx3cr1^+^ cells expressing mCherry were observed at the two vs. 7-8 week timepoint in blood (**Fig. 6G**; p=0.0175) and spleens (**Fig. 6H**; p=0.0012). Not only do we confirm stable expression of mCherry in Cx3cr1^+^ cells in the brain between both timepoints, but we also show a clear turnover of peripheral cell transgene expression 7-8 weeks post-TAM. Notably, basal expression of mCherry in all tissues was detected in vehicle and TAM treated animals, closely matching our IHC assessments above.

**Figure 6.**
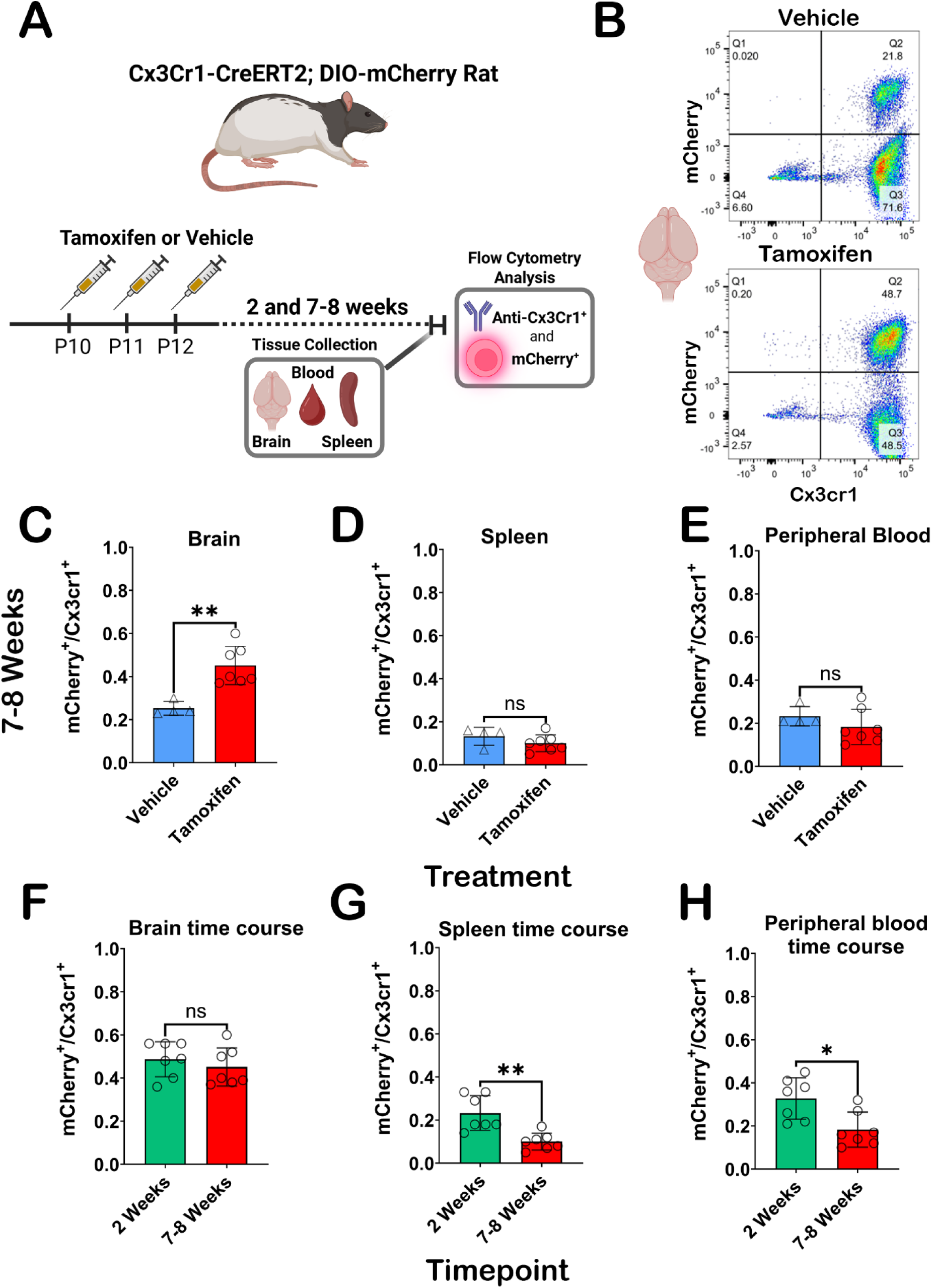
Flow cytometry analysis of Cx3cr1-CreERT2^+/-^; DIO-mCherry^+/-^ rat brain, blood and spleen. **(A)** Experimental overview for flow cytometry experiments. **(B)** Representative flow cytometry detection of ectopic mCherry and Cx3cr1 (BV604) antibody results following cell gating, doublet exclusion, and live/dead staining from the brain in vehicle (top) and TAM (bottom) at 7-8 weeks. (**C-D**) Graphical representations of mCherry expression in Cx3cr1^+^ cells at 7-8 weeks in brain (**C)**, spleen (**D**), and peripheral blood (**E**) in vehicle or TAM treated animals. Significant elevation in mCherry expressing Cx3cr1^+^ cells was only observed in the brain (p=0.0061) at this timepoint. mCherry expressing Cx3cr1^+^ cells in tissue from TAM treated Cx3cr1-CreERT2^+/-^ ; DIO-mCherry^+/-^ rats is stable from 2-8 weeks in the brain (**F**), while there are significant decreases in mCherry expressing Cx3cr1^+^ cells in peripheral blood (**G**) and spleens (**H)** in the 2 vs. 7–8-week groups (p=0.0175 and p=0.0012, respectively). Statistical analysis was performed using the non-parametric two-tailed Mann Whitney test (ns=not significant;*p<0.05; **p<0.01).

## 3. Discussion

The studies above describe the generation and characterization of a novel transgenic rat containing a microglia/monocyte-specific Cre-driver that is tamoxifen inducible: the Cx3cr1-CreERT2 rat. The development of this rat adds to the breadth of literature using Cx3cr1-promoter driven transgene expression models already available in mice. In tandem, we also further characterize and utilize a Cre-dependent fluorescent protein reporter rat, DIO-mCherry, described previously (Bäck, Necarsulmer, et al., 2019). We advance our understanding of the DIO-mCherry rat by demonstrating transgene expression in response to AAV-mediated Cre delivery and breeding this rat with the Cx3cr1-CreERT2 rat to enable inducible mCherry transgene expression in microglia. As more transgenic rats become available, tools for specifically manipulating specific cell populations, such as microglia, will become routine and invaluable for understanding niche cell contributions to pathology and behavior.

In the absence of Cre recombinase, the DIO-mCherry rats do not produce mCherry fluorescence in the brain, but following an AAV-iCre injection, we readily observe mCherry fluorescence in Cre expressing cells—primarily in neurons due to the specificity of the AAV vector used. The Cx3cr1-CreERT2^+/-^; DIO- mCherry^+/-^ rats have detectable mCherry^+^ cells in various brain regions in the absence of tamoxifen treatment; however, in TAM treated rats, the number of Iba1^+^mCherry^+^ cells increases 2.7-4.8-fold. When using Cx3cr1-CreERT2 mice to examine microglia response to gene disruption/expression, it is common to have a “washout period” of three weeks after TAM treatments to allow for recombined Cx3cr1^+^ peripheral cell turnover and replacement by non-recombined progenitor cells. Although this strategy does eliminate most peripheral transgene expression, some is still observed even past the expected turnover rate. Turnover of Cx3cr1^+^ peripheral circulating and immune cells occurs in approximately 3-4 weeks (Goldmann et al., 2013; Parkhurst et al., 2013; van Furth & Cohn, 1968; Yona et al., 2013), while self-renewing microglia (Ajami et al., 2007) persist for a much longer, 3–4-month period (Askew et al., 2017). This fast turnover rate of peripheral cells and slow turnover of microglia in the brain allows for the use of Cx3cr1-Cre animals to successfully target microglia without confounding peripheral recombination (Goldmann et al., 2016).

Concerns of Cx3cr1-CreER leakiness in mice — Cre or transgene expression in cells other than microglia in the brain— have sparked numerous studies evaluating the specificity (Bedolla et al., 2023; Zhao et al., 2019) and off target effects of this model in mice (Chappell-Maor et al., 2020; Sahasrabuddhe & Ghosh, 2022). In the Cx3cr1-CreERT2^+/-^; DIO-mCherry^+/-^ rats, our experiments show minimal non-microglial cells expressing the mCherry transgene following VEH or TAM treatment, and some TAM independent expression of mCherry in the spleen, blood, brain, and primary microglial cultures. To examine tim- and cell-specific expression of mCherry in the periphery and central nervous system, we utilized flow cytometry (FC).

In these double hemizygous rats, detection of mCherry in the absence of TAM is not unexpected, as CreERT2 is known to have some basal level of Cre recombinase activity in the absence of ligand, as demonstrated in the Cx3cr1-CreERT2 transgenic mouse by several groups (Bedolla et al., 2024). In the periphery, Cx3cr1 is expressed in several cell types in the blood and spleen, including leukocytes that populate the red pulp and marginal zone of the spleen(Bedolla et al., 2024; Zhao et al., 2019). While we demonstrate that mCherry transgene is stably expressed from 2-8 weeks in the brain, peripheral cell expression is elevated at two weeks post-TAM treatment, with significant decreases by eight weeks. Like other studies examining transgene expression in Cx3cr1-CreER mice (Swirski et al., 2009), we also observe low level peripheral expression of mCherry in blood and spleen. In Cx3cr1-CreER mice, tamoxifen independent transgene expression was detectable up to 9-10 weeks. Although we observe mCherry expression in a small number of Iba1^-^ cells in the brain post-TAM treatment, cells are primarily Iba1^+^ indicating that the Cre expression is limited to microglia. (Bedolla et al., 2024).

Given that the intended uses for the Cx3Cr1-CreERT2 rats are to study the role of microglia in rat cognition and behavior, we tested the transgenic rats for potential deficits in cognitive tasks. We utilized sensory preconditioning, which permits assessing typical reward conditioning (Pavlov, 2010) - the bell means food is coming – in addition to inference (Brogden, 1939), the person who rings the bell also predicts food. While reward learning is computationally simple and easily predicted by learning models, inference as assessed by sensory preconditioning is not (Miller et al., 1995). Critically, both processes were intact in transgenic rats and functionally identical to wild type, which is not always the case in transgenic lines (Costa et al., 2021). Inference is a critical component of mammalian cognition (Jones et al., 2012), and intact inference, in addition to intact reward-learning, would indicate that the Cx3cr1-CreERT2 transgene did not generally interfere with learning processes. Our behavioral data indicate that Cx3Cr1-CreERT2^+/-^ rats are capable of both simple and complex cognitive operations and will therefore provide a suitable model for their assessment in a broad range of models and conditions with impaired or altered microglial function.

Previous work showed that when crossed with DAT-Cre rat, mCherry expression was primarily localized to the DAT-expressing cells of the rat midbrain (Bäck, Necarsulmer, et al., 2019). Here we show that AAV-Cre can be used in DIO-mCherry rats to produce mCherry in rat brain which we observed mostly in neurons. This was expected based on the AAV serotype and promoter used primarily expresses transgenes in neurons when injected into the rat STR. In addition to experiments delivering AAV-iCre to label cells with mCherry, there is a growing list of Cre-driver rats available commercially and through repositories which could be crossed with DIO-mCherry to label cells throughout the body. Here we demonstrate that the mCherry expression is sufficiently expressed for flow cytometry analysis of brain and blood.

Microglia are the resident immune cells of the CNS, but that is not the only important function they hold. They play a key role in many different processes across the CNS and present different characteristics across different brain regions (De Biase et al., 2017). The diverse context-dependent phenotypes and region-specific differences of microglia functions (De Biase et al., 2017) presents many new research questions that can be tackled using genetic models such as our Cx3cr1-CreERT2 rat.

Rat models are valuable tools in research, especially in neuroscience, where their larger brain size and more complex behaviors offer advantages over mouse models. The development of a new rat model capable of expressing tamoxifen-inducible transgenes in CX3CR1-expressing cells *in vivo* opens new opportunities to study immune cells throughout the body, as well as microglia in the CNS. This advancement is particularly significant for investigating neuroinflammation and exploring complex behaviors and cognitive processes.

## 4. Materials and Methods

### 4.1 Animals

All animal experiments were conducted on adult rats in accordance with the National Institute of Health (NIH) guidelines for animal research. All experiments were approved by the Animal Care and Use Committees (ACUC) of the Intramural Research Programs at the National Institute on Drug Abuse and the National Institute of Mental Health. The rats were group-housed in a 12 h light-dark cycle with *ad libitum* access to rodent chow and water. Animals involved in behavior studies utilized an altered food schedule described below. The assignment of animals to experimental groups was based on genotyping results (for most experiments) and based on parental genotypes and *post hoc* genotyping of subjects (i.e. primary microglial cultures).

The Cx3cr1-CreERT2 transgenic animals were generated at Cyagen US Inc. in specific pathogen free facilities that have been AAALAC accredited and OLAW assured.

The DIO-mCherry transgenic animals were generated at the NIMH Transgenic Animal Core. A transgene cassette expressing Cre-dependent mCherry from the human EF1α promoter was liberated from the viral vector backbone by digestion with MluI and RsrII restriction enzymes. The DNA fragment was gel purified and injected into the pronuclei of LE rat zygotes. These were transferred to pseudo pregnant LE females as single cell zygotes on the day of injection or allowed to develop to 2-cell embryos the next day before transfer. The resulting line, LE-Tg(DIO-mCherry)2Ottc, is registered at the Rat Genome Database (RGD: #8693598) and deposited at the Rat Resource and Research Center (RRRC #687; University of Missouri, Columbia MO, USA). Animals were bred with WT LE rats from Charles River Laboratories.

### 4.2 AAV Vector construction

#### pAAV EF1α iCre

The open-reading-frame encoding iCre (based on pBSII-iCre (Shimshek et al., 2002), was amplified by PCR with linkered primers and inserted the BamHI and EcoRI sites of pOTTC293 (Addgene; #60057) using ligation-independent cloning (In-Fusion; Clontech). The resulting plasmid (pOTTC556, Addgene; #210416) was grown in Stbl3 cells (Invitrogen) and verified by sequencing before packaging into AAV particles.

#### pAAV-EF1α CreERT2

The coding region for CreERT2 was amplified from MSCV CreERT2 puro (a gift from Tyler Jacks; Addgene; #22776) and inserted into an AAV packaging plasmid flanked by the EF1α promoter and the poly-adenylation signal from bovine growth hormone using ligation-independent cloning (In-Fusion, Clontech). The resulting plasmid (Addgene; #210415) was grown in Stbl3 cells (Invitrogen) and verified by sequencing before being used as a backbone for the recombineering template.

### 4.3 BAC Recombineering Cx3cr1-CreERT2

A bacterial artificial chromosome (BAC) containing the rat Cx3cr1 gene (CH230-274J7) was obtained from the Children’s Hospital Oakland Research Institute (CHORI), and transformed into the recombineering strain (GS1783, DH10B λ *cI*857 Δ(*cro-bioA*)<>*araC*-P_BAD_ *I-sceI*). “En passant mutagenesis” (detailed below) was used to replace the start codon of Cx3cr1 with a cassette containing the CreERT2 coding region and the polyadenylation signal from bovine growth hormone (Tischer et al., 2010).

The pAAV EF1α CreERT2 plasmid was digested with PshAI to allow the insertion of an I-SceI restriction site and a kanamycin selectable marker, which was amplified from pEP-KanS [a gift from Nikolaus Osterrieder (Addgene; #41017)]. A 50 bp portion of the CreERT2 coding region (spanning amino acids 278 to 293) was duplicated on both sides of the insertion to provide regions of homology that facilitate intramolecular recombination and marker removal in subsequent steps. The final plasmid, pAAV EF1α CreERT2 (KanS) was used as a template to amplify the entire CreERT2-(I-SceI-KanS) cassette by PCR with primers containing 5’ linkers containing 50 bp of homology with the sequence adjacent to the Cx3cr1 start codon. This PCR product was electroporated into heat shocked GS1783 cells containing the Cx3cr1 BAC, and the transformants were selected on LB agar plates containing chloramphenicol and kanamycin. Successful transformants were screened by PCR across the 5’ and 3’ junctions and sequence verified for error-free insertion. The kanamycin resistance marker was removed and the CreERT2 coding region restored by the simultaneous induction of I-SceI with arabinose and the induction of lambda-red genes by heat shock. Cultures were diluted and streaked onto plates containing LB agar plates containing chloramphenicol. Successful recombinants were screened by PCR and verified by sequencing and pulse-field agarose gel electrophoresis.

The final CX3CR1-CreERT2 BAC was prepped using NucleoBond Xtra BAC columns (Macherey-Nagel) and used by Cyagen US Inc. to create the transgenic rat line.

### 4.4 Creation of Cx3cr1-CreERT2

For the Cx3cr1-CreERT2 rat, the modified bacterial artificial chromosome (BAC) containing the rat Cx3xcr1 gene with the CreERT2 coding region was injected into the embryos of LE rats by Cyagen US Inc. and the resulting progeny were screened for the transgene using PCR based genotyping. The resulting line, LE-Tg(CX3CR1-CreERT2)3Ottc, is registered at the Rat Genome Database (RGD: 13441557) and deposited at the Rat Resource and Research Center (RRRC; #858; University of Missouri, Columbia, MO). Animals were bred with WT LE rats from Charles River Laboratories for 10 generations before crossing with DIO-mCherry rats for experiments.

The DIO-mCherry rat was previously described (Bäck, Necarsulmer, et al., 2019). Briefly, the transgene expression cassette was isolated from the plasmid pAAV EF1α DIO mCherry [pOTTC337, Addgene; #47636 (Richie et al., 2017)] by digestion with MluI and RsrII and agarose gel excision using Macherey-Nagel Gel and PCR cleanup kit. This purified DNA fragment was microinjected into fertilized oocytes harvested from a LE rat by the NIMH Transgenic Core. Surviving pups were screened for the integrated transgene by PCR genotyping. This line (LE-Tg(DIO-mCherry)2Ottc) has been registered with the Rat Genome Database (RGD; #8693598) and deposited at the Rat Resource and Research Center (RRRC; #687). Herein, ‘‘LE-Tg(DIO-mCherry)2Ottc’’ rats are referred to as ‘‘DIO-mCherry’’ rats. Animals were bred with WT LE rats from Charles River Laboratories for 4–5 generations, prior to being crossed with Cx3cr1-CreERT2 animals for detection of mCherry expression.

### 4.5 Genotyping

Genomic DNA was isolated from tissue biopsies or primary cells using a Macherey-Nagel Tissue Spin kit. PCR genotyping was performed using gene-specific primers and OneTaq polymerase (New England Biolabs) with a 68°C annealing temperature and indicated extension time in a Bio-Rad T100 thermocycler. Primers used are described in **Table 2**. Amplification products were analyzed by agarose gel electrophoresis or by fragment analysis on an AATI 5200 Fragment Analyzer (Agilent).

**Table 2.**
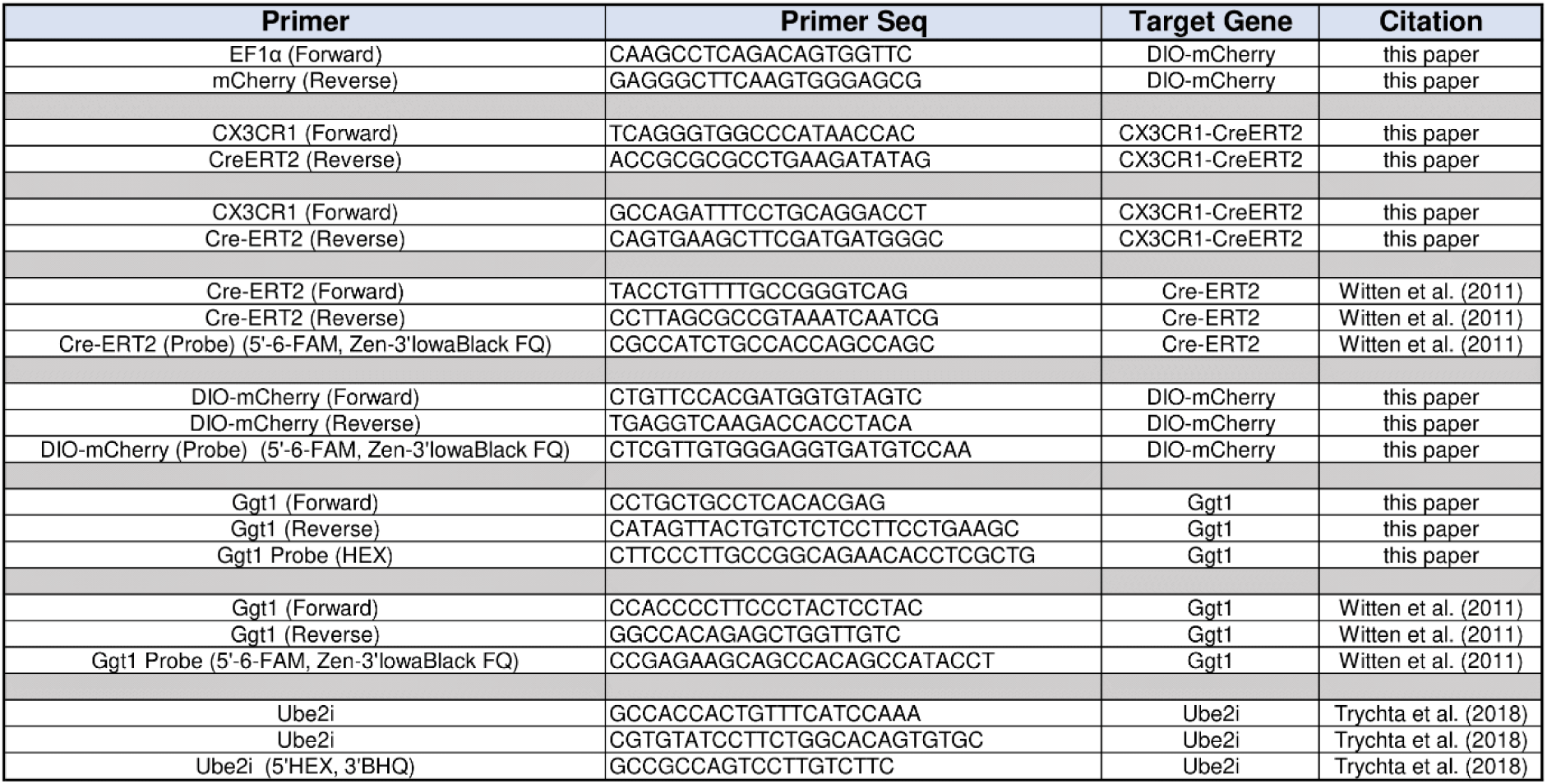
Primers and Probes.

### 4.6 Copy number Analysis

Genomic DNA was isolated as described above and served as template for droplet digital PCR (ddPCR) using ddPCR Supermix for Probes (No dUTP) (Bio-Rad; #1863024) or QIAcuity Probe PCR Kit (Qiagen; # 250102), respectively and FAM- or HEX-labeled probe assays for CreERT2, DIO-mCherry, and rat Ggt1 (located on chromosome 20) [**Table 1**]. For ddPCR, droplets were generated and analyzed using a QX200 AutoDG Droplet Digital PCR System and amplification reactions were run in a Bio-Rad T100 thermocycler. For dPCR, QIAcuity Four Platform System (Qiagen) with QIAcuity Nanoplates 8.5k (Qiagen; #250011) were used.

### 4.7 RNA isolation, cDNA synthesis, and dPCR

RNA was isolated from brain tissue using the RNeasy Lipid Tissue kit (Qiagen, #78404) with an additional DNAse treatment (RNAse-Free DNAse set; Qiagen, #79254). Polytron PT1200 was used to mechanically homogenize tissue samples. RNA quality was analyzed using Bioanalyzer 2100 (Agilent, #G2939B) and RNA 6000 Nano Reagent Kit (Agilent, #50671511) with RNA Nano Chip (Agilent, #50671511) according to Bioanalyzer instructions. cDNA was made using the iScript cDNA Synthesis kit (BioRad, #1708891) according to manufacturer’s instructions. 500 ng or RNA was used as input for the generation of cDNA. cDNA reaction was diluted 2.5 times and 5uL of cDNA were used per dPCR reaction.

Reaction mix consisting of 0.8uM primers, 0.4uM probes, and 4X Probe Mix (Qiagen, #250102) were pipetted into the QiAcuity 8.5k 24 well Nanoplate (Qiagen, #250011) and run on QiAcuity Four Platform dPCR system (Qiagen, #911042). Samples were primed according to the QIAGEN standard priming profile, then cycled once at 95°C for 2’, followed by 40 cycles of 95°C for 15s and 60C for 30°s in sequence. Exposure of both the green and yellow channels were set at 500ms with a gain of 6. Cx3xr1 and Cre copy numbers were normalized to Ube2i. Primers and probes used for these assays are listed in **Table 2**.

### 4.8 AAV Injections

Adult LE rats were anesthetized using intraperitoneal (i.p.) injections of ketamine (80mg/kg) and xylazine (8mg/kg) and placed into a stereotaxic frame (Stoelting Co., Wood Dale, IL). Rats were given intracranial injections of AAV-EF1α-iCre into the STR using the following coordinates: AP +0.5; ML ±3.0, DV -6.0 ± 4.0 mm from bregma. 2.0 µL of AAV [8.90E^10^ vg/mL] was injected into both hemispheres at a rate of 0.5µL/min using a Nanofil 10 μL syringe with a 33G blunt needle and a UMP4 microinjector pump (World Precision Instruments, Sarasota, FL).

### 4.9 Tamoxifen injections

Ten days after pups are born, Tamoxifen (TAM) (Millipore Sigma #T5648-1G) was administered at a dose of 60mg/kg through IP injections once a day over a course of three days. TAM was dissolved in sunflower oil + 10% ethanol. Solution was heated to 37°C, sonicated, and degassed until fully dissolved. Animals were perfused two, four, six, or eight weeks after the final injection day.

### 4.10 Tissue Processing and Imaging

#### 4.10.1 Perfusions

Animals were perfused with 0.9% saline followed by 4% freshly prepared paraformaldehyde (PFA). Brains were removed and placed into 50 ml conical tube with 4% PFA for 2 hours, 4°C. After 2 hours, the PFA was removed and cold 18% sucrose in phosphate buffer (1xPB) was added to the conical tube and returned to 4°C overnight. The following day, after achieving saturation, 18% sucrose solution was removed and cold 30% sucrose in 1xPB was added and returned to 4°C overnight. After saturation, brains were rapidly frozen of a bed of dry ice. After fully frozen brains were stored at -80°C.

#### 4.9.2 Sectioning

Brains were sectioned (30 µm) using a Leica CM1950 cryostat and stored in cryopreserve solution (20% glycerol, 2% DMSO in 1XPB). A hole was punched into the right hemisphere during sectioning to aid in consistent orientation of slices during imaging. Sectioning was completed in a sequential series of 3 sections through the 3 different brain regions: PFC/STR (Bregma 2.20 to 1.70 mm), HC (Bregma -3.14 to -4.16 mm), MB (Bregma -4.80 to -6.04 mm). Brain coordinates were determined using online tool by Matt Gaidica developed with Paxinos and Watson (2006) rat brain atlas. Sections were stored at -80°C.

#### 4.9.3 Immunohistochemistry

Sections were thawed to room temperature and washed in PB solution on a shaker at room temperature for 10 minutes twice. After the third wash, sections were transferred to blocking solution (4% Normal Goat Serum in 1xPB solution with 0.3% Triton X-100) for 1 hr at room temperature on a shaker. After blocking, sections were transferred to primary antibodies, mouse anti-mCherry (Living Colors; #632543; Lot# 2206232A and #2406A38A; 1:1500 dilution) and rabbit anti-Iba1 (Wako; #019-19741; Lot #PAM4131 and #LKJ2979; 1:1500 dilution) in blocking solution and placed on a shaker at 4°C for overnight incubation. Following overnight incubation, sections were transferred to PB solution for a 3 x 10 min wash on a room temperature plate shaker. The sections were then transferred to blocking solution with secondary antibodies goat anti-mouse (488) and goat anti-rabbit (680) at dilutions of 1:1500 (Invitrogen; #A21109 and #2831347, respectively). The plate was covered with aluminum foil and placed on a shaker at room temperature for 2 hours. The sections were then transferred to 1xPB solution for a 10 min wash at room temperature. Sections were then transferred to blocking solution with DAPI (ThermoFisher; #D3571) at a 1:3000 dilution for 10 minutes. Then sections were transferred to 1xPB for a final wash step. Sections were then mounted on a charged glass slide and covered with a glass cover slip with Mowiol (SigmaMillipore; #475904-100GM) mounting medium.

#### 4.9.4 Confocal specifications, imaging parameters, and analysis parameters

A Nikon Eclipse NI standing upright confocal microscope was used for imaging. Images were acquired using the 405 (DAPI), 488 (anti-mCherry), 561 (mCherry), and 640 (anti-Iba1) lasers. Laser power and gain was set to 7 and 90 respectively for the 405 channel, 5 and 105 for the 488, 5 and 115 for the 561, and 7 and 90 for the 640 channels. These specifications were established using a brain of known genotype (positive control). All users were blinded to rat genotype during the imaging process. Six z-stacks of 10 steps at 1 µm increments were taken per region of interest (PFC, STR, HC, and MB). Z-stacks were taken at 20X magnification and 1024 1024 resolution. One z-stack was taken per brain hemisphere for PFC, STR, and MB over 3 sections for six unique z-stacks per region per animal. Z-stacks were taken of the dentate gyrus and CA3 region on the same hemisphere for HC sections over 3 hippocampi for six unique z-stacks per HC per animal. Approximate location of imaging on coronal section is shown in **Supplemental Fig. 3**. Maximum intensity projections were generated from all channels of each z-stack via *Fiji* v1.54 (Schindelin et al., 2012). This maximum intensity image was used for quantitative data collection. Brightness and contrast were enhanced separately and uniformly across all samples in the 640 channel (Iba1) via *Fiji* v1.54, reducing background noise within the z-stacks. Brightness and contrast settings were set to a pixel value of 1500-1510 for 16-bit (.nd2 file) max projection. Two blinded scorers counted all the maximum projection images for the following: mCherry positive cells, anti-mCherry positive cells and anti-Iba1 positive cells. A record of mCherry^+^/Iba^-^ cells was kept for each image. Counts were averaged across six images for each brain region per animal (n= 6-7 animals per group). Iba1^+^ soma immunoreactivity was used a minimum qualifier for a cell to be counted as a microglial cell. For statistical analysis of imaging data, one-way or two-way ANOVA with Tukey’s multiple comparisons test was run for all data sets. *p*-values < 0.05 were considered statistically significant (\**p* < 0.05; \*\**p* < 0.01; \*\*\**p* < 0.001). All statistical analyses were conducted in GraphPad Prism (version 10.1.2). Overlap coefficient was determined using JACoP: Just another Co-localization Plugin (Bolte & Cordelieres, 2006) for *Fiji*.

### 4.11 Behavior

#### 4.11.1 Subjects

Thirty-three adult male (n=15) and female (n=18) wild type (n=18) and Cx3cr1-CreERT2 transgenic (n=15) LE rats weighing between 270- and 320g during the time of experiments were pair housed and given ad libitum food and water except during behavioral training and testing. Four rats (3 female wild type, 1 female transgenic) failed to show typical reward conditioning, defined as higher responding to the first presentation of cue B than to the first presentation of cue D during the B-D reward conditioning probe test. These rats were therefore excluded from all analyses, leading to the following final group sizes: male WT (n=7), male transgenic (n=8), female WT (n=8), female transgenic (n=6). The effect of cue during the critical A-C probe test did not depend on sex or genotype (sex main: F_(1,25)_=3.04, *p*=0.09; genotype main: F_(1,25)_=0.87, *p*=0.36 sex × cue: F_(1,25)_=3.67, *p*=0.07; sex × cue × genotype interaction: F_(1,25)_=0.74, *p*=0.40). Five days prior to behavioral training rats were food restricted to 12g (males) or 10g (females) of standard rat chow per day, following each session. Rats were maintained on a 12 hr/12 hr light/dark cycle and tested during the light phase between 8:00 am and 2:00 pm five days per week.

#### 4.11.2 Apparatus

All chambers and cue devices were commercially available equipment (Coulbourn Instruments, Allentown, PA). A recessed food port was placed in the center of the right wall of each chamber and was attached to a pellet dispenser that dispensed three 45 mg banana flavored sucrose pellets (Bioserv; #F0024, Flemington, NJ) during each rewarded cue presentation. Auditory cues consisted of a pure tone, siren, 2 Hz clicker, and white noise, each calibrated to 70 dB. Cues A and C were white noise or clicker; cues B and D were tone or siren; all cues were counterbalanced across rats, sexes, and groups such that every possible permutation of cue identities, sex, and group was equally present to the extent possible.

#### 4.11.3 Behavioral training

Procedures generally followed previous work (Hart et al., 2022; Hart et al., 2020; Jones et al., 2012; Sharpe et al., 2017).

*Preconditioning.* Five days following the beginning of food restriction, rats were shaped to retrieve pellets from the food port in one session. Twenty pellets were delivered on a 3-6m variable interval schedule. Rats then underwent two days of preconditioning. During each session rats received trials in which two pairs of auditory cues (A→B and C→D) were presented in blocks of six trials. Block order was counterbalanced across rats and sessions. Cues were each 10 s long, and pairs were delivered on a 3-6m variable interval. Behavioral responses reported here and following are the session-average percentage of time spent during each 10s cue in the food port, minus the preceding 10s (CS-baseline).

*Conditioning.* Following preconditioning rats received 6 days of reward conditioning, 1 session per day. During each session rats received six trials of cue B paired with pellet delivery and six trials of D paired with no reward. Three pellets were dispensed during cue B at 3, 6.5, and 9 s into the 10s presentation. Cue D was presented for 10 s without reward. Cues were presented in random order on a 3-6m variable interval.

*Inference probe test.* Following conditioning, rats were given a probe test during which they received non-reinforced presentations of cues A and C to test for inference behavior. Cues A and C were presented 6 times each in blocks of 3 trials with 3-6m between cue presentations. The order of the cues/blocks was counterbalanced across subjects.

*Reward conditioning probe test.* Following the A-C probe and on a separate day, rats were given a probe test during which they received non-reinforced presentations of cues B and D to test for typical reward conditioning behavior. Cues B and D were presented 6 times each in blocks of 3 trials with 3-6m between cue presentations. The order of the cues/blocks was counterbalanced across subjects.

*Histology.* Rats were sacrificed by isoflurane overdose and perfused with PBS. Half of the rats were then perfused with 4% formaldehyde in PBS. Following perfusion, brains were extracted and flash frozen (PBS perfusion) or post fixed in 4% formaldehyde (PBS-formaldehyde perfusion).

#### 4.11.5 Statistical analyses

Data were collected using Graphic State 4 software (Coulbourn Instruments, Allentown, PA). Raw data were processed and analyzed in Matlab 2023a (Mathworks, Natick, MA) to extract percentage of time spent in the food port during and during the 10s preceding cue presentation. Analyses were conducted on responding during cue presentation, minus this 10s baseline period. We compared responding during preconditioning and probe testing by 3-way (sex, genotype, cue) ANOVA. To assess whether rats acquired higher responding to the reward-paired cue during conditioning, we subtracted D responding from B (B-D) and compared this measure across the six conditioning sessions using 3-way ANOVA (sex, genotype, session). Analyses were run on session-averages.

### 4.12 Flow cytometry and organ isolation

#### 4.11.1 Blood and organ collection

Animals were deeply anesthetized in a drop jar saturated with isoflurane. The thoracic cavity was opened, and the right atrium of the heart was punctured. Prior to perfusion, 100 µL of blood was collected from the right atrium and mixed in a 1:2 MACS buffer (1xPBS pH 7.4, 0.5% BSA, 2 mM EDTA) on ice. Animals were perfused with 0.9% saline prepared with 1:1000 heparin (v/v). Spleen and brain was removed and placed into DMEM High Glucose on ice.

#### 4.11.2 Tissue processing for flow cytometry

Each brain was individually hemisected and minced on ice, followed by enzymatic digestion in RPMI (Gibco; #11875-101) with 1.25 mg/ml ColD (Sigma; #11088882001) and 50 ug/ml DNase I (Sigma; #DN25-100MG) for 30 min at 37°C using the ADBK_01 program in C-tubes (Miltenyi; #130-093-237). Tissue was then triturated and strained through a 70 μm pre-wetted filter and subjected to a 30% and 70% isotonic Percoll (Cytiva; #17089101) gradient centrifugation for 30 min at 1,000*g* at 4°C. The interphase layer between the Percoll gradients was collected and used for downstream flow cytometry analysis.

Spleens were individually crushed using a syringe plunger in MACS buffer (1xPBS pH 7.4, 0.5% BSA, 2 mM EDTA). Blood was mixed with MACS buffer, as described above, to prevent clotting. Spleens and blood were treated with ACK (Ammonium-Chloride-Potassium) lysing buffer at RT for 10 min to selectively remove red blood cells.

#### 4.11.3 Flow cytometry

Single cell suspensions from each tissue were stained for 20 min in FC block (Biolegend; #156604; 1:100) in MACS buffer on ice. Cells were centrifuged at 300g at 4°C for 5 minutes and resuspended in PBS plus Zombie Aqua Live-Dead stain (Biolegend; #77143; 1:1000) and incubated on ice for 20 minutes. Cells were centrifuged at 300g at 4°C for 5 minutes and resuspended in MACS buffer with anti-Cx3cr1 (Brilliant Violet 605; BioLegend; #149027; 1:200) and incubated on ice for 20 minutes. Cells were then washed two times in ice cold MACS and strained for flow cytometry analysis. Flow samples were analyzed using a Becton Dickinson LSRFortessa equipped with 405, 488, 561, and 647 nm lasers. Forward Scatter (FSC) and Side Scatter (SSC) were respectively used to identify cells of interest and eliminate doublet cells for gating analysis. A third gate using Zombie Aqua staining allowed for identification of live cells subsequently analyzed for ectopic mCherry expression (using PE-CF594 detection) and Cx3cr1 expression (using BV605 detection). Gates were established using WT controls and single stains. Gating strategy for all tissues is shown in **Fig. S8 A-C**, with WT (negative controls) and Cx3cr1-CreERT2; DIO-mCherry gates for stained brain, spleen, and peripheral blood are shown in **Fig. S9 A-C**. Percent of Cx3cr1^+^ cells that also express mCherry^+^ are reported within the manuscript.

### 4.12 Primary microglia isolation and maintenance

Microglia isolation follows method described in (Sepulveda-Diaz et al., 2016) with the following changes: T-75 flasks were coated with 10 ml of PEI diluted 1:500 in borate buffer. Flasks were placed in 37⁰C, 5.5% CO2 incubator for 1 hr. After 1 hr flasks were washed for a total of 3x with 1X PBS. Pups were collected on day 2 postnatal from DIO-mCherry and Cx3cr1-CreERT2 crossing. Whole brains were isolated and placed individually in a 5 mL tube containing 2 ml of DMEM. Tail snips were collected and sent to Transnetyx for genotyping. Using a P1000 pipet the brain was mechanically dissociated by trituration. Once a homogenized suspension was reached debris were allowed to settle for 2 minutes and supernatant passed through a 100 µm cell strainer into a pre-cooled 50 mL tube. An additional 2 mL of complete media (DMEM, 10% HI-FBS, 1%P/S) was added to the tube containing debris and trituration was repeated. Again, debris was allowed to settle and supernatant collected into the tube already containing the first collection.

Combined supernatant was spun down at 1000rpm, 4°C, for 5 min. Supernatant was discarded and cell pellet resuspended in 5 mL of complete media. Cell suspension was added to the PEI coated T-75 flask containing 5 mL of complete media. 10 ml of media total per flask. After 72 hrs, 8 mL of media was removed, and 18 mL of fresh complete media was added.

At 18 days in vitro (DIV18) cells were detached from the flask using 0.25% trypsin. The Countess (Invitrogen) was used to count viable cells that were plated on PEI coated 24- and 96- well plates at densities of 1.5 x 10^5^ cells / well and 3 x 10^4^ cells/ well, respectively. DIV19 cells are treated with 1 µM 4-OHT. At DIV22 cells were washed once with PBS and fixed with 4% PFA for 1 hr at room temperature.

After fixation, cells were permeabilized with 0.1% Triton, 0.2% BSA for 15 min at RT then blocked for 1 hr at room temperature in PBS containing 0.1% Triton, 2% BSA, and 5% goat serum. Cells were incubated overnight with anti-Iba1(Wako:1:500) at 4°C. Following overnight incubation cells were washed 3× with PBS then secondary rabbit antibody. Alexa fluor 488 (Invitrogen) was added for 1 hr at room temperature. Cells were washed 3× with PBS and nuclei stained with DAPI (Invitrogen; 1:1000) for 10 min at room temperature.

### 4.12 Primary cortical neuron isolation, maintenance, and treatment

Following methods previously described in (Bäck, Dossat, et al., 2019) primary cortical neurons (PCNs) were isolated from E15 Sprague-Dawley rat embryos and plated onto PEI coated 96 well plates, DIV0. On DIV6 cells were transduced with AAV1-EF1α-CreERT2-bGHpA +/- AAV1-DIO-Cherry. 100µL of media was removed from each well and 5µL of each virus was added for 2 hrs. On DIV8 cells were treated with 1 μM 4-OHT by 50% media exchange. Cells were imaged for DAPI and red fluorescence from DIV9 to DIV15.

## Acknowledgements

This work was supported by the Intramural Research Programs at the National Institute on Drug Abuse and National Institute on Aging as well as the NIDA HIV Research Program and NIH Office of AIDS Research. We thank Mark Jackson (animal husbandry), Tearra McGlotten (animal husbandry), Sarah Blossom (animal husbandry), Yajun Zhang (genotyping) and Nick Edwards (vector construction) for their technical contributions. AAV viral vectors were packaged by Doug Howard in the NIDA Genetic Engineering and Viral Vector Core (formerly the Optogenetics and Transgenic Technology Core).

---

**Figure S1:**
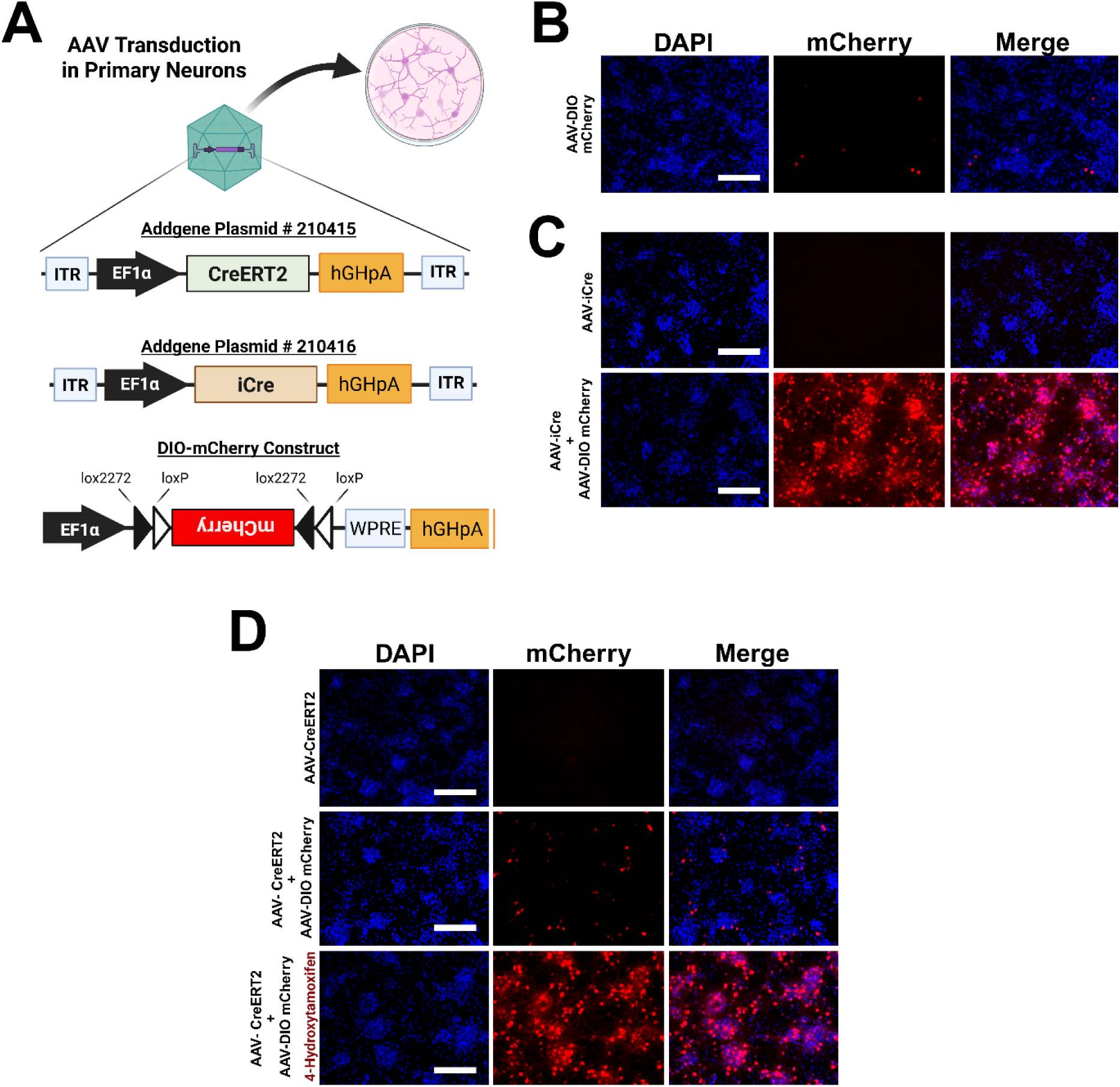
Verification of CreERT2 and DIO-mCherry activity prior to generating rat. **(A)** Primary cortical neurons were used to test our CreERT2 (Addgene: #210415) and DIO-mCherry constructs delivered using AAV transductions. **(B)** AAV DIO-mCherry produces minimal Cre-independent mCherry expression in PCNs. **(C)** Co-delivery of AAV-iCre (Addgene: #210416) and AAV DIO-mCherry to PCNs induces robust mCherry expression in PCNs. (D) AAV-CreERT2 and AAV DIO-mCherry co-transduction induces minimal tamoxifen-independent mCherry expression in PCNs, while addition of 4-hydroxytamoxifen (1µM) following the same AAV-transduction generates robust mCherry expression. (Scale Bar: 200 µm).

**Figure S2:**
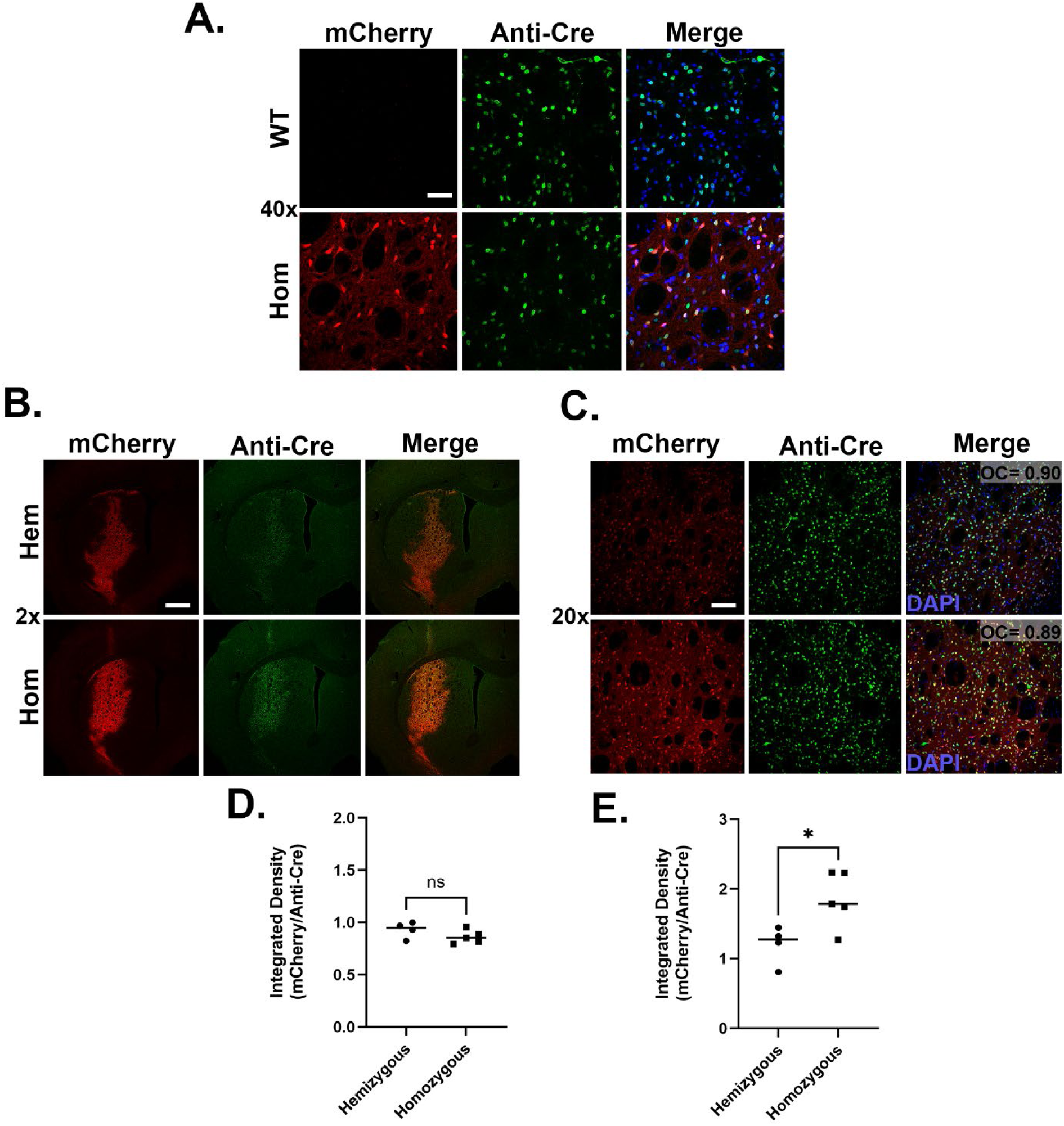
Hemizygous vs. Homozygous LE DIO-mCherry rat characterization. (**A**) Representative 40x magnification images from both Long Evans (LE) wildtype (WT) and DIO-mCherry (DIO-mCherry^+/+^) homozygous animals two weeks after delivery of AAV-Cre into the striatum. In the presence of the AAV delivered Cre, mCherry expression is absent the WT animal and prominent in the DIO-mCherry animal. The mCherry signal is colocalized with Cre immunoreactivity (scale bar: 100 μm). (**B**) Representative low magnification images from hemizygous (Hem) and homozygous (Hom) DIO-mCherry animals 2 weeks after injections with AAV-Cre. Colocalization between Cre immunoreactivity and mCherry epifluorescence is evident in both animals (scale bar: 1 mm). (**C**) Higher magnification images from same brain sections to demonstrate colocalization of the two signals (scalebar= 200 µm). Quantification of integrated density ratios of Cre immunoreactivity/DAPI fluorescent signal shows comparable expression of Cre in both Hem and Hom animals (**D**). The integrated density ratio of mCherry/Cre immunoreactivity is significantly higher in the Hom animals compared to Hem animals. (**E**). Each data point represents average integrated density of two coronal sections/animal (n=4-5 per group). Unpaired t-test was used for statistical analysis (*p<0.05).

**Figure S3:**
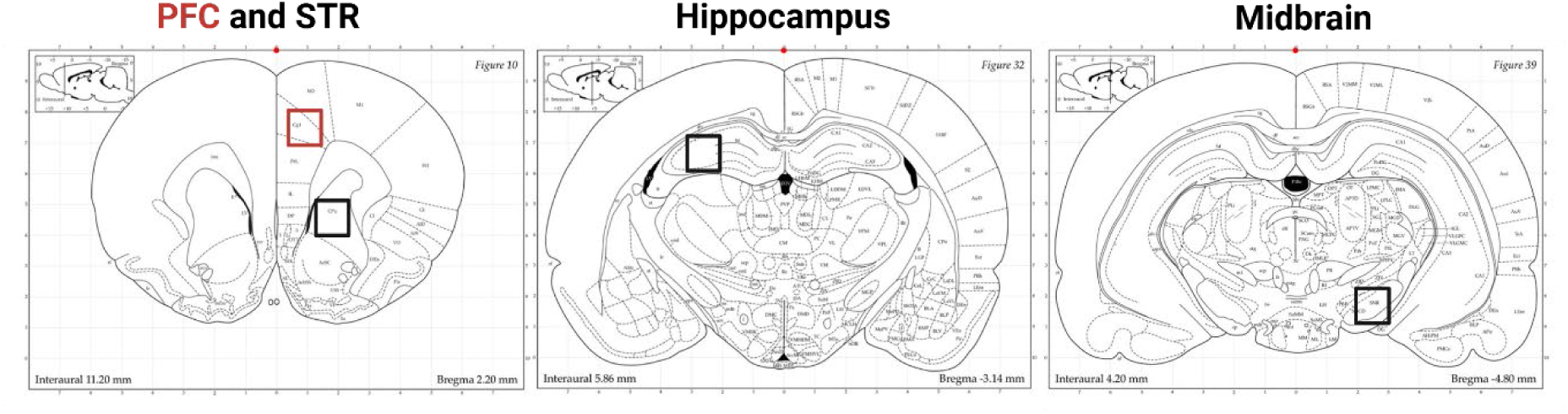
Brain regions where mCherry expression was analyzed for colocalization with the Iba1. Images were acquired from prefrontal cortex (PFC), striatum (STR), hippocampus (HC) and midbrain (MB). Coronal schematics were modified from Paxinos and Watson (2006).

**Figure S4:**
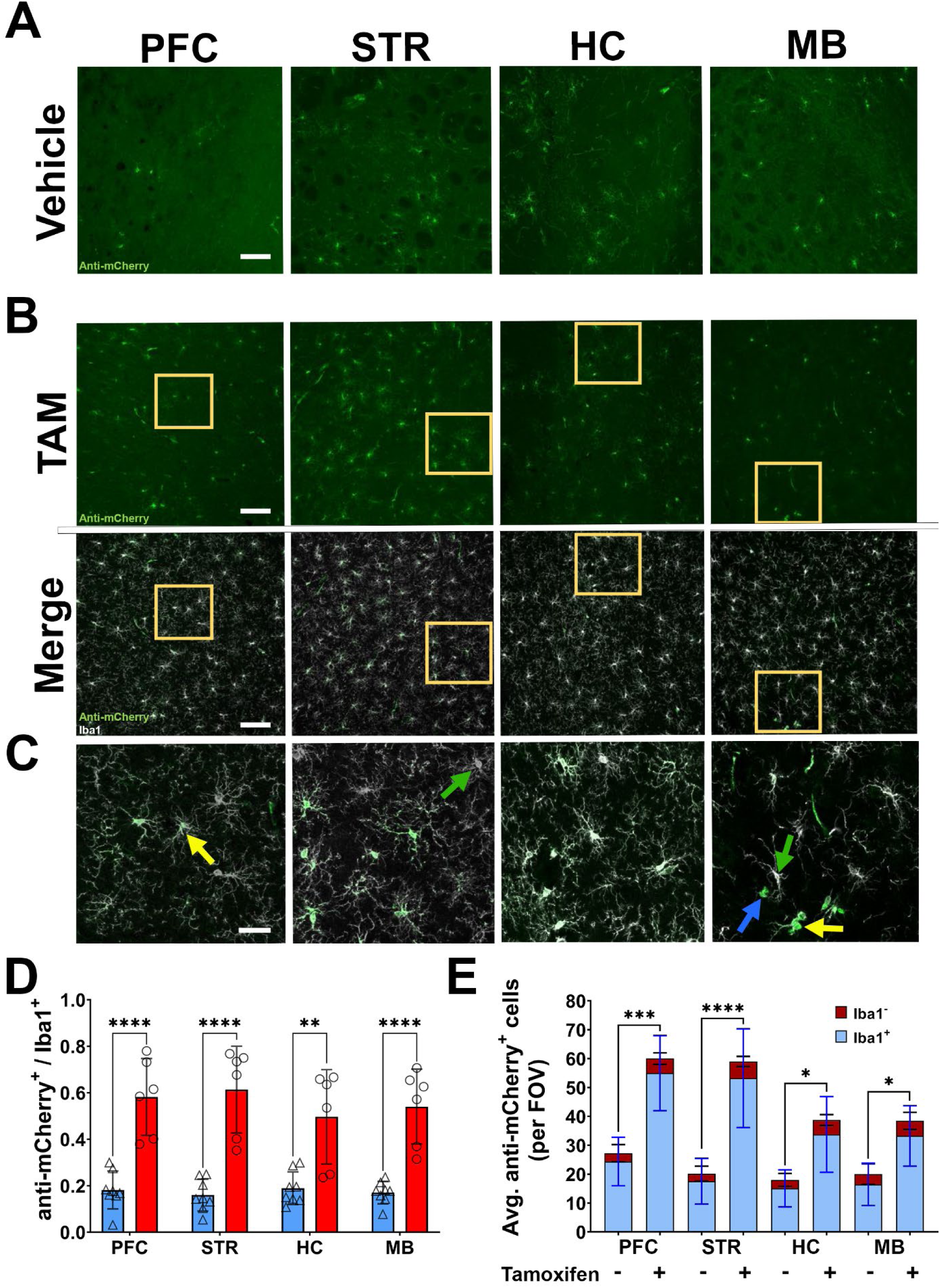
Microglial Cre activity confirmation in LE Cx3cr1-CreERT2^+/-^; DIO-mCherry^+/-^ rat using anti-mCherry antibody. Male and female Cx3cr1-CreERT2^+/-^ ; DIO-mCherry^+/-^ rats received three i.p. injections of either tamoxifen (TAM) (male, n=3; female, n=4) or vehicle (male, n=4; female, n= 3) at P10. Brain tissue was collected 8 weeks after injections. Brains were sectioned and images taken proximal to the prefrontal cortex (PFC), striatum (STR), hippocampus (HC), and midbrain (MB) sections. Representative regional images show anti-mCherry (green) expression in vehicle **(A)** and TAM **(B)** injected rats (scale bar:100 µm) (top panel of **B**). Colocalization of anti-mCherry (green) and Iba1 (white) (Merge) in TAM injected rats is shown in the Merge panel of **(B)**. Regions shown at higher magnification in **(C)** (scale bar= 30 μm) are highlighted in yellow boxes from **(C)**. Yellow arrows indicate representative Iba1^+^ anti-mCherry^+^ cells; green arrows highlight Iba1^+^ anti-mCherry^-^ cells; blue arrows show Iba1^-^ anti-mCherry^+^ cells. **(D)** Ratio of anti**-**mCherry^+^ /Iba1^+^ cells in vehicle (n=7) and tamoxifen (n=6) treated rats (male and female animals combined). **(E)** Raw counts of mCherry^+^ cells in vehicle and tamoxifen treated animals. Each bar indicates average quantity of anti-mCherry^+^ cells that are Iba1^+^ (blue) and Iba1^-^ (red) per field of view. Two-way ANOVA with Tukey’s multiple comparison with S.D. error bars (**p<0.01; ***p<0.001; ****p<.0001).

**Figure S5:**
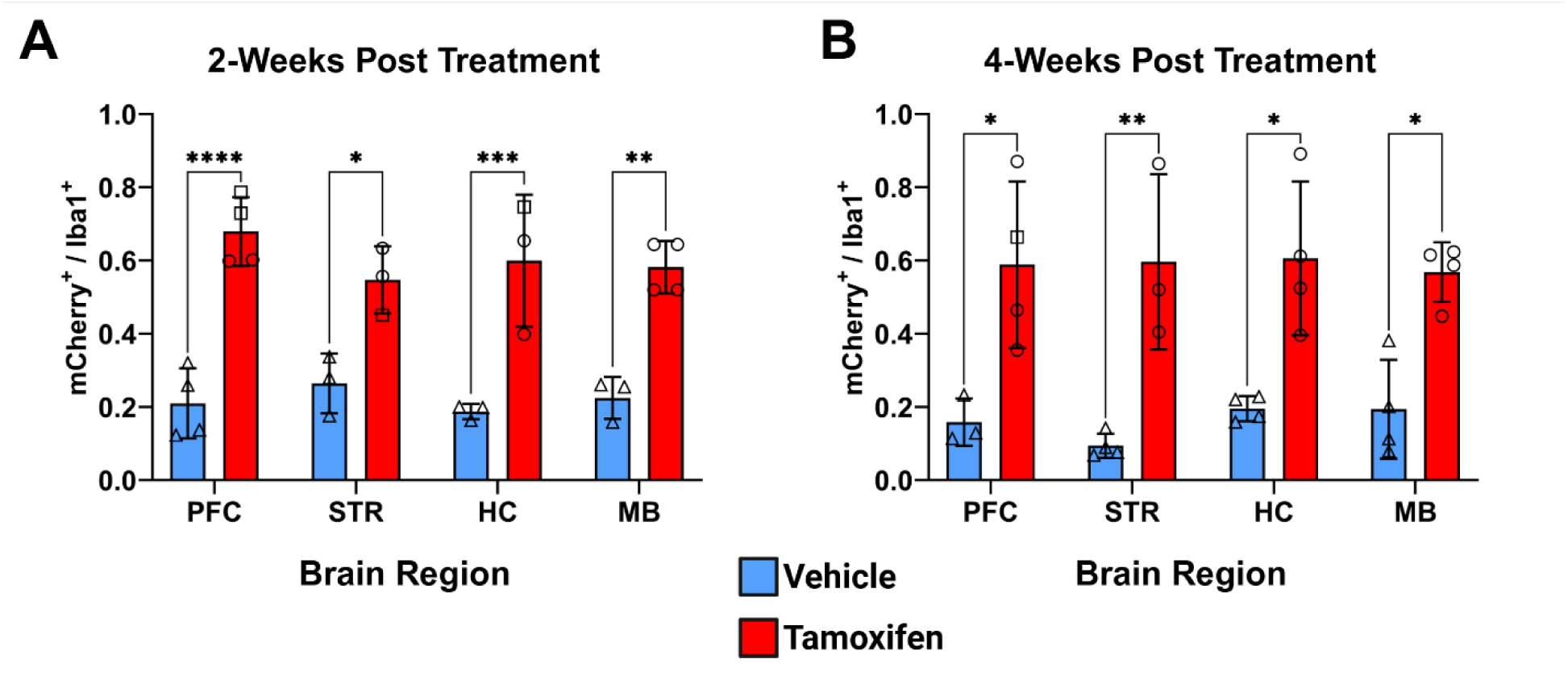
Two- and four-week post tamoxifen mCherry^+^/Iba1^+^ cell count ratios in the LE Cx3cr1-CreERT2^+/-^; DIO-mCherry^+/-^ rat brain. Male and female double transgenic animals (Cx3cr1-CreERT2**^+/-^**; DIO-mCherry**^+/-^**) received three-five i.p. injections of either tamoxifen (TAM) (male, n=3; female, n=4) or vehicle (male, n=4; female, n= 3) at P10. Brain tissue was collected 2- or 4- weeks after vehicle or tamoxifen injections. Brains were sectioned and images taken proximal to the prefrontal cortex (PFC), striatum (STR), hippocampus (HC), and midbrain (MB). Square symbols indicate animal was given five TAM injections. Two-way ANOVA with Tukey’s multiple comparison (*p<0.05; **p<0.01; ***p<0.001; ****p<.0001) with error bars indicating S.D.

**Figure S6:**
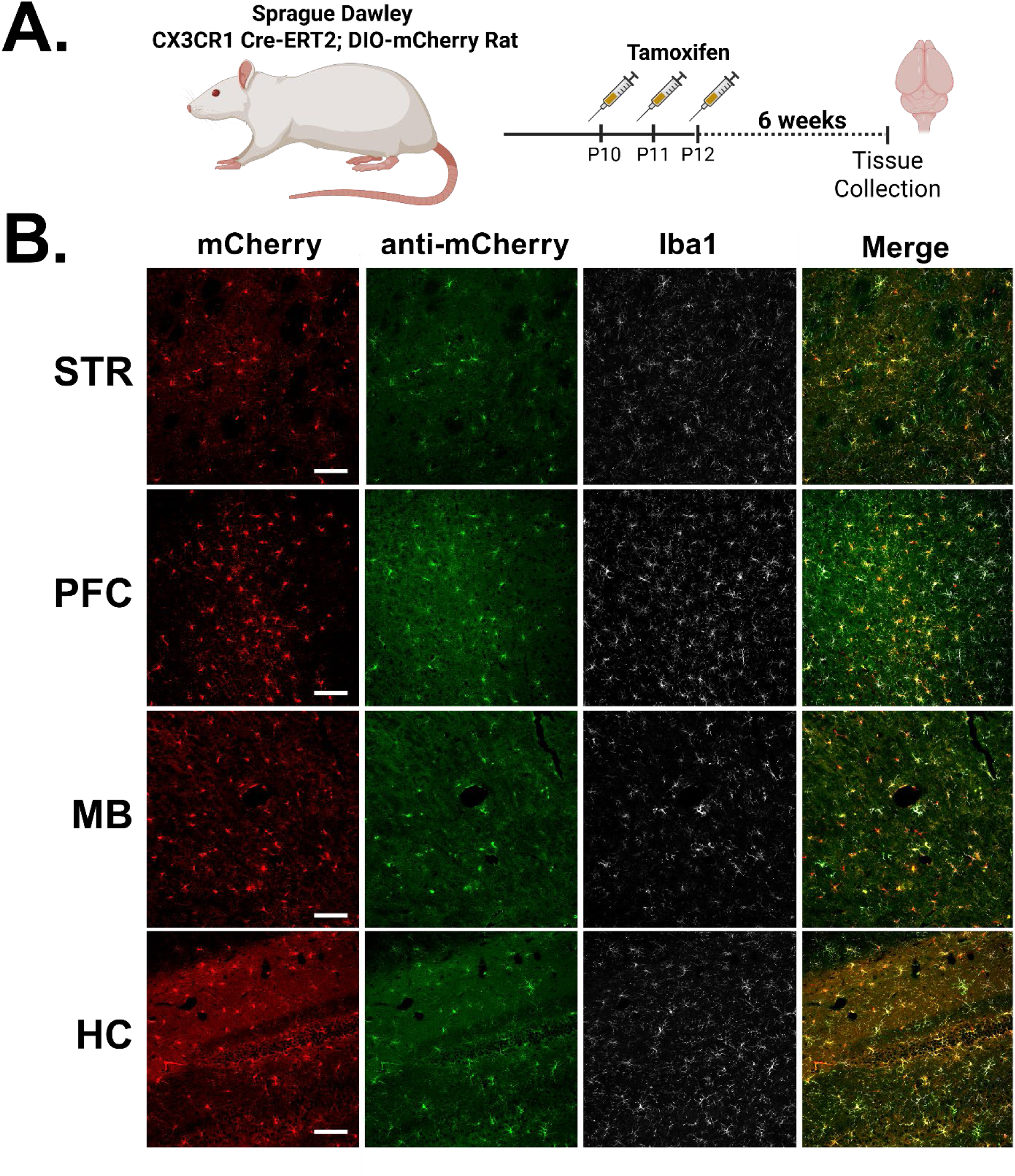
Specificity of microglial Cre activity in SD background Cx3cr1- CreERT2^+/-^; DIO-mCherry rat^+/-^. **(A)** Sprague Dawley Cx3cr1-CreERT2; DIO-mCherry rats were injected with TAM (60mg/kg) once daily for three days. Brains were collected 6 weeks post TAM, sectioned and analyzed for ectopic mCherry expression (red) and anti-Iba1 (white) and anti-mCherry (green) immunoreactivity in the prefrontal cortex (PFC), striatum (STR), hippocampus (HC), and midbrain (MB) **(B)**. Colocalization of all three channels visualized in the “Merge” column. Scale Bar =100 µM

**Figure S7:**
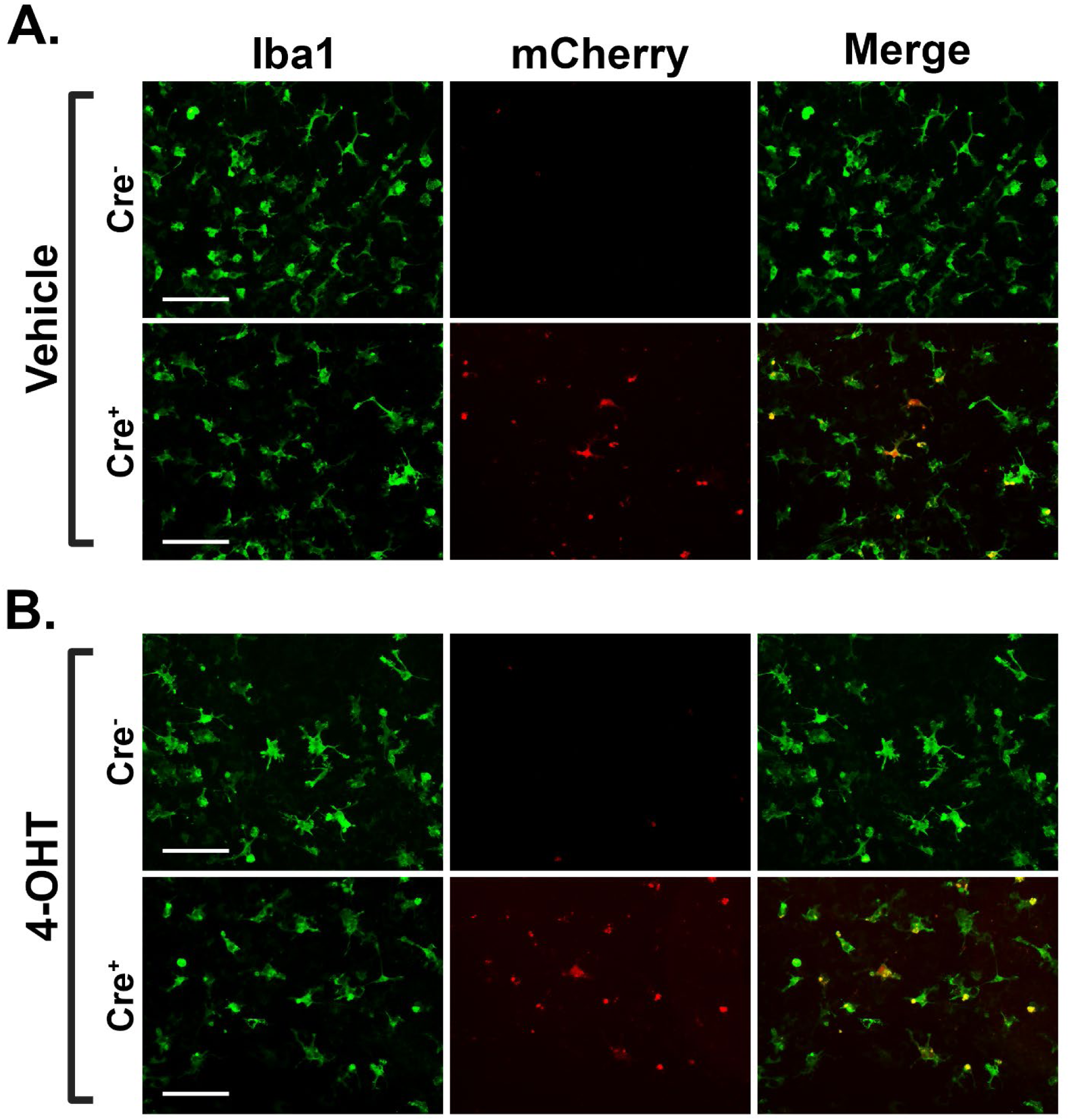
Primary Microglia from SD background Cx3cr1-CreERT2^+/-^; DIO mCherry^+/-^ animals. Primary microglia were isolated and plated from 2 days post-natal pups of Cx3cr1-CreERT2^+/-^ × DIO-mCherry^+/-^ crossing. Genotyping was performed from postmortem tissue collection. Cells were treated with either Vehicle (**A**) or 4-hydroxytamoxifen (4-OHT) (**B**). mCherry expression was not detected in microglia from animals without the CreERT2 (Cre^-^) transgene in either the Vehicle or 4-OHT treatment groups. Vehicle treated microglia with the CreERT2 (Cre^+^) show mCherry expression, indicative of basal levels of active Cre (**A**). mCherry signal is increased in Cre^+^ microglia treated with 4-OHT (**B**).

**Figure S8:**
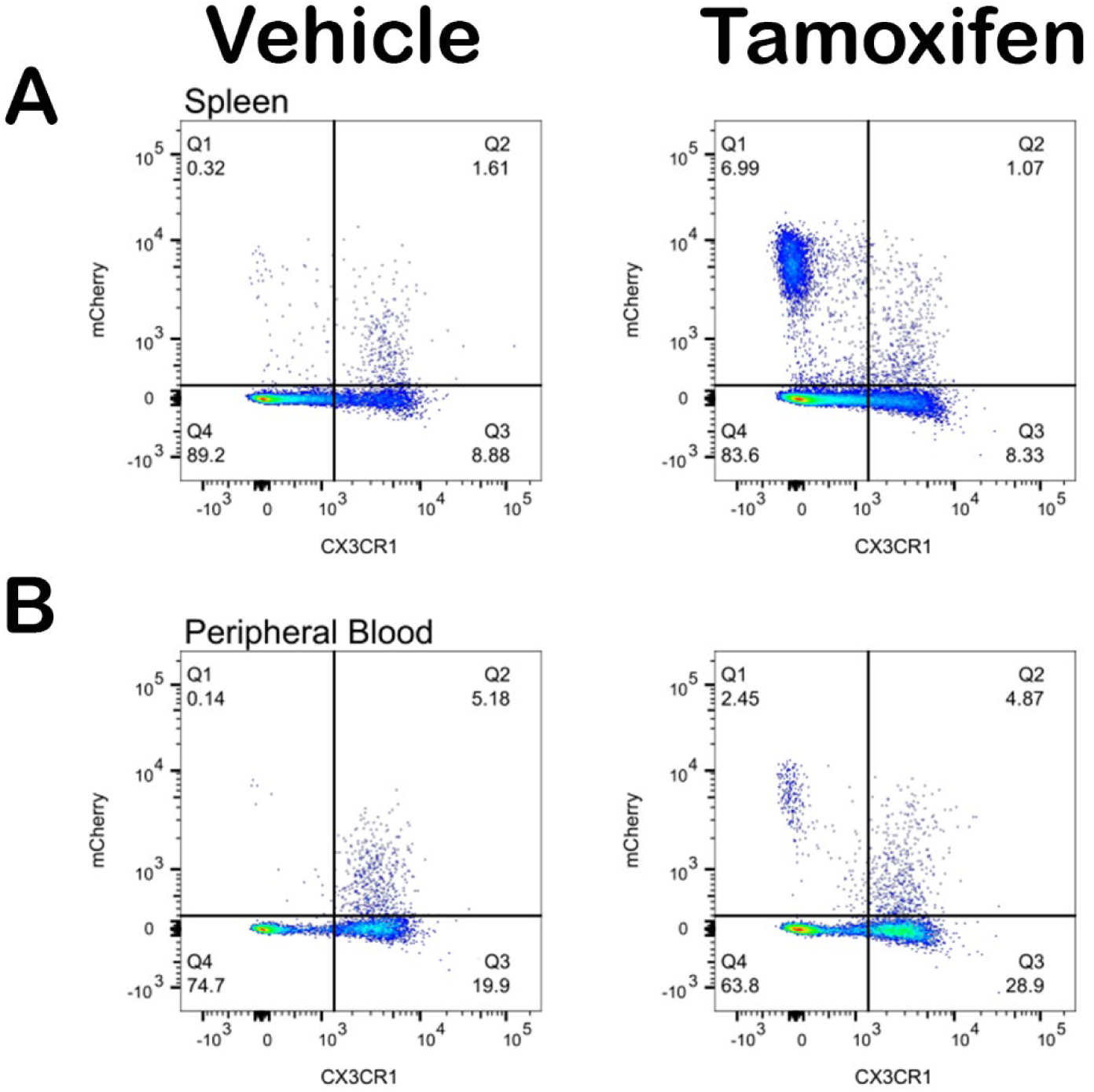
Flow cytometry gating comparisons for spleen and peripheral blood. Splenocytes (**A**) and peripheral blood (**B**) from either vehicle (left) or TAM (right) treated Cx3cr1- CreERT2^+/-^; DIO-mCherry^+/-^ were analyzed via flow cytometry. Representative gating charts are shown.

**Figure S9:**
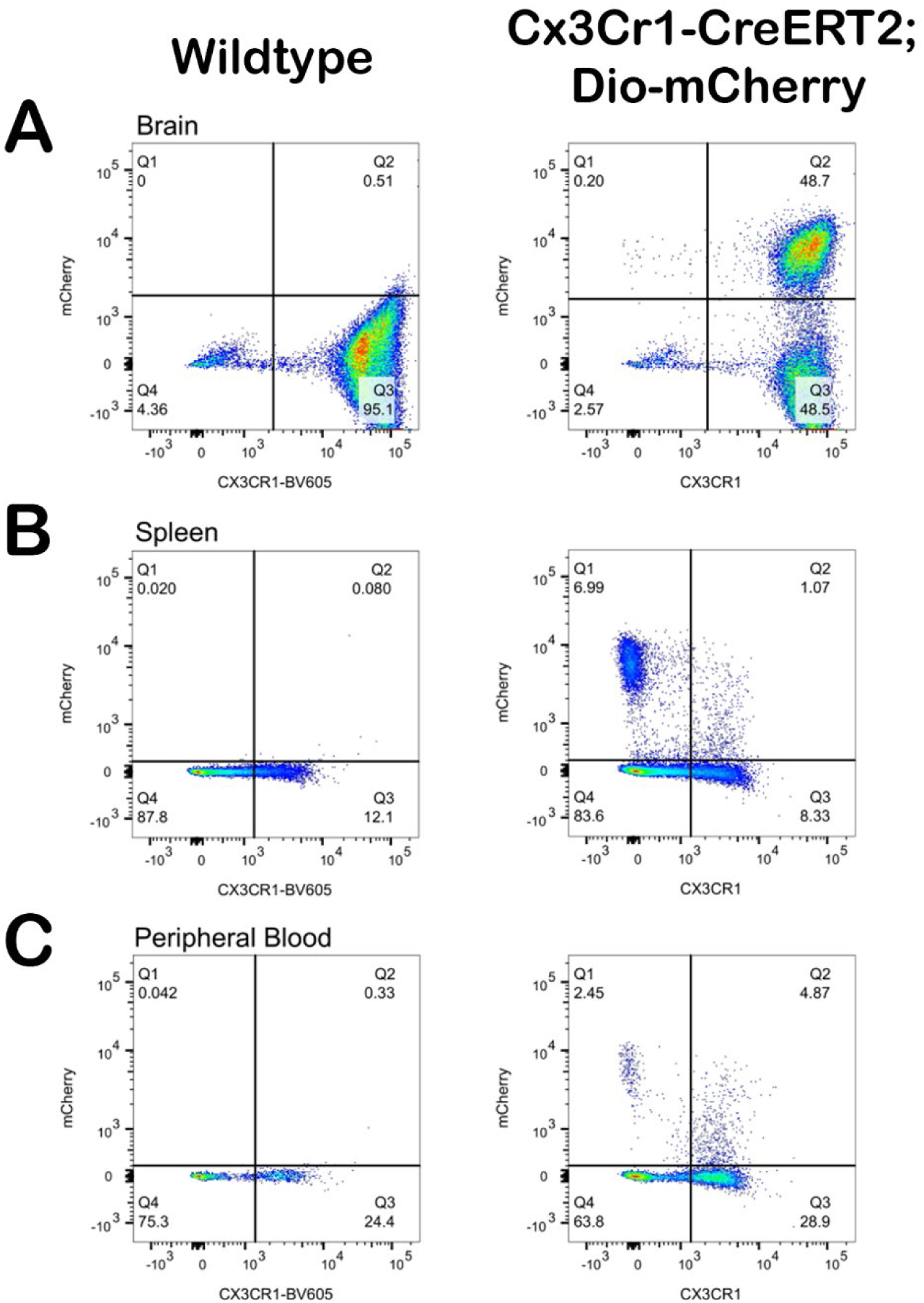
Flow cytometry gating determination of brain, spleen, and peripheral blood via stained WT negative controls. TAM treated WT animals were used to gate thresholds for mCherry expression in brain (**A**), spleen (**B**), and peripheral blood (**C)** of TAM treated CreERT2^+/-^; DIO-mCherry^+/-^ rats.

